# Management of Hsp90-Dependent Protein Folding by Small Molecule Targeting the Aha1 Co-Chaperone

**DOI:** 10.1101/412908

**Authors:** Jay Singh, Bradley Tait, Naihsuan Guy, Jeffrey C. Sivils, David Culbertson, Chad Dickey, Szu Yu Kuo, Jason E. Gestwicki, Darren M. Hutt, Jane H. Dyson, Marc B. Cox, William E. Balch

**Affiliations:** The Scripps Research Institute, Department of Molecular Medicine, Skaggs Institute of Chemical Biology, La Jolla, CA; Brad Tait Enterprise, 80 Christian Way North Andover, MA 01845; Department of Biological Sciences and Border Biomedical Research Center, University of Texas at El Paso, El Paso, TX; Department of Integrative Structural and Computational Biology, The Scripps Research Institute, La Jolla, CA; Department of Molecular Medicine and Byrd Alzheimer’s Research Institute, University of South Florida, Tampa, FL, USA; Department of Pharmaceutical Chemistry, University of California San Francisco, San Francisco, CA 94158

## Abstract

The core cytosolic Hsp90 chaperone/co-chaperone complex plays a critical role in proteostasis management of human health and disease. To identify novel compounds that alter the ability of the Hsp90 co-chaperone Aha1 to modulate the ATPase activity found in multiple folding diseases ranging from steroid hormone receptor (SHR) sensitive prostate cancer to tauopathies associated with neurodegenerative diseases, we employed a <u>h</u>igh <u>t</u>hroughput <u>s</u>creening (HTS) assay to monitor selectively <u>A</u>ha1-<u>s</u>timulated <u>H</u>sp90 (ASH) ATPase activity. The ASH assay identified SEW04784 (SEW), a small molecule that disrupts ASH activity without inhibiting the basal Hsp90 ATPase activity. NMR analysis reveals that SEW binds to the C-terminal domain of Aha1 to disrupt its asymmetric binding to Hsp90 leading to abrogation of its chaperoning activity of Hsp90. SEW exhibits therapeutic potential by blocking the transcriptional activity of prostate cancer (PCa) associated variants of the androgen receptor (AR) in a cell-based model of PCa. Additionally, SEW exhibits the ability to clear toxic, phosphorylated tau aggregated species associated with tauopathies. By not directly impacting the basal ATPase function of the abundant and ubiquitous Hsp90, SEW could provide a therapeutic approach for mitigation of client-specific proteostatic disease.

## Introduction

Heat shock protein 90 (Hsp90), is one of the most abundant proteins in the cell, reaching an expression level of 2-5% of total protein. It is evolutionarily conserved, ubiquitously expressed and plays an essential role in cell signaling, proliferation, and survival^1-11^ as part of the proteostasis protein folding management system^8,9,12-14^. Hsp90 is an ATP-dependent molecular chaperone which, together with its regulatory co-chaperones and the Hsp70 chaperone/co-chaperone system, form the centerpiece of cellular proteostasis management of protein folding^8,12,15^. The Hsp90 chaperone and its multiple co-chaperones regulate the stability, trafficking, function and degradation of a wide range of client proteins and is therefore central to all cellular processes. Their specific roles in protein folding biology largely remains an enigma. Moreover, the value of Hsp90 and its co-chaperones as therapeutic targets for management of proteostasis-dependent responses to familial and somatic genetic disease, and/or physical stress-related disease that impacts the function of the protein fold remains to be demonstrated in the clinic. Hsp90 plays a critical role in management of Spatial Covariance (SCV), a new biological principle based on Gaussian regression to describe central dogma and its role in dictating the biology of the protein fold in response to evolvability and natural selection^16^.

In humans, the genes that code for Hsp90 isoforms^5^ include the mitochondrial tumor necrosis factor (TNF) receptor associated protein (TRAP1), the luminal endoplasmic reticulum localized, glucose regulated protein 94 (GRP94) (HSP90β), the stress inducible Hsp90α (HSP90AA) and the constitutively expressed Hsp90β (HSP90AB)^5,14^. The functional Hsp90 protein in cells exists as a homodimer, with each protomer composed of 4 domains: an N-terminal domain (NTD) containing the ATP-binding pocket, a charged linker connecting the NTD and the middle domain (MD), the MD, which is involved in client and co-chaperone binding as well as ATP hydrolysis and a C-terminal domain (CTD) involved in dimerization and recruitment of TPR containing co-chaperones via its MEEVD motif^5,14^. The Hsp90 dimer undergoes a series of structural rearrangements during its ATP-binding and hydrolysis cycle. In the nucleotide-free (apo) form, the C-terminally dimerized Hsp90 adopts an open conformation, which closes upon ATP binding and hydrolysis, a process that promotes protein folding and is assisted by the action of its multiple co-chaperones^5,14^, including the accelerator of Hsp90 ATPase (Aha1)^17-32^.

Aha1 is a two domain, Hsp90 co-chaperone conserved from yeast to man^21,23,33-35^. It competes with other Hsp90 co-chaperones for binding to Hsp90 and contributes to the functional activation of client proteins, including kinases, steroid hormone receptors and transcription factors ^1,4,5,33,36,37^, by stimulating the ATPase activity of Hsp90^38-40^. We have previously shown that Aha1 binding to Hsp90 is asymmetric. It bridges the middle domain of one of the Hsp90 protomers in the Hsp90 dimer with the N-terminal domain of the other Hsp90 molecule to promote the N-terminal dimerization of Hsp90 and thereby stimulate its ATPase activity^38-40^. The silencing of Aha1 decreases client protein activation and increases cellular sensitivity to Hsp90 inhibitors^38,41,42^. In cystic fibrosis (CF), a reduction of the Aha1 expression levels *in viv*o corrects the trafficking and functional defect associated with the deletion of Phe508 (F508del) variant of the <u>c</u>ystic fibrosis transmembrane <u>c</u>onductance regulator (CFTR), the most abundant CF-associated mutation^32^. These data suggest an important role for reduced ATPase activity in the stabilization of mutant CFTR, a hypothesis that we posited can be extended to other Hsp90-associated diseases including cancer and neurodegenerative diseases^32,38,42^.

Numerous small molecule Hsp90 inhibitors have been developed and tested in clinical trials for cancer which exhibit an “addiction” to Hsp90^18,43-53^. Many of these compounds, including the natural products geldanamycin (GA) and radicicol and their derivatives^54-57^, target the NTD of Hsp90, blocking its ATPase activity by competing for the ATP binding pocket. To date, none of these compounds have met with clinical success in cancer indications^58-62^, an effect that is likely due to the mechanism of action of these compounds, which are likely to completely inhibit the general activity of the essential Hsp90 protein responsible for maintaining proteostasis balance^8^ in the cell. To circumvent this issue, new Hsp90 inhibitors have been developed to target co-chaperone binding. These include novobiocin and other compounds^51,63-69^ that competes for binding to the C-terminal domain of Hsp90 and blocks the binding of TPR containing co-chaperones, and the recently-identified small molecules that disrupt the binding of Aha1 by directly interacting with Hsp90^17,20^ as well as compounds indirectly inferred to interact only with Aha1^18^. The potential for small molecule regulators of Hsp90 co-chaperone sensitive disease remains to be established in the clinic.

To identify novel compounds that alter the ability of Aha1 to modulate the ATPase activity of Hsp90, we employed a <u>h</u>igh throughput <u>s</u>creening (HTS) which monitors the ATPase activity of Hsp90 activity in the presence of Aha1. Using this <u>A</u>ha1-<u>s</u>timulated <u>H</u>sp90 (ASH) assay we identified SEW04784 (SEW), a novel scaffold that disrupts ASH activity without inhibiting the basal Hsp90 ATPase activity. Using NMR, we show that SEW binds to the C-terminal domain of Aha1 to disrupt its asymmetric binding to Hsp90 leading to abrogation of the chaperoning activity of Hsp90 *in vitro*. *In vivo*, SEW exhibits therapeutic potential by blocking the transcriptional activity of prostate cancer (PCa) associated variants of the androgen receptor (AR) in a cell-based model of PCa. Additionally, SEW exhibits the ability to specifically clear toxic, phosphorylated tau species associated with tauopathies, such as Alzheimer’s disease. SEW is predicted to have high value in the clinic given the client-dependent role of Aha1 in mitigation of human protein misfolding diseases without directly impacting the basal ATPase function of the abundant and ubiquitous Hsp90.

## Results

### Small molecule inhibitors of Aha1-stimulated Hsp90 (ASH) ATPase activity

To identify small molecules that can inhibit ASH activity, we adapted an assay that is based on the ability of colored products to quench the inherent fluorescence of white microtiter plates^70,71^. In brief, inorganic phosphate (P_i_), originating from the Hsp90-mediated hydrolysis of ATP, can be complexed with molybdate to form phosphomolybdate. The latter can then bind to the quinaldine red (QR) dye to form a colored product that quenches the inherent fluorescence of the white microtiter plates (**Figure 1A**). In our study, compounds that inhibit the ATPase activity of Hsp90 will lead to a lower production of P_i_ and exhibit less fluorescence quenching.

**Figure 1.**
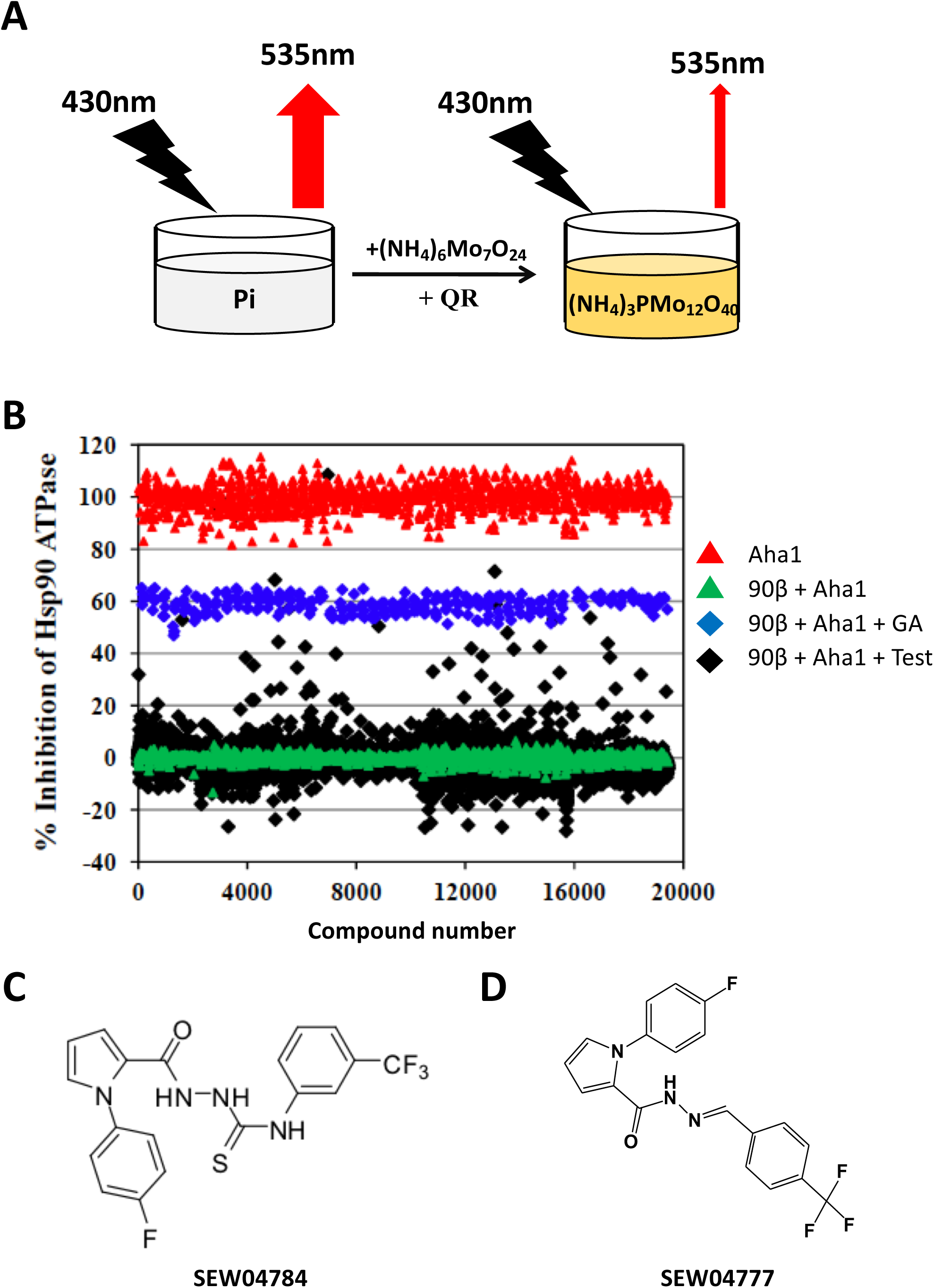
High throughput screen identifies small molecule inhibitors of the Aha1-stimulated Hsp90 ATPase (ASH) assay. **A.** Schematic diagram depicting the principle of the quinaldine red-based quenching of the inherent fluorescence of white microtiter plates. **B.** Scatterplot of the high throughput screen to identify inhibitors of the ASH activity from the Maybridge discovery library of chemical compounds. The data is presented as a percent inhibition of the ASH activity, where Hsp90+Aha1 is set as 0% inhibition (green triangles) and Aha1 is set as 100% inhibition (red triangles). The positive control for Hsp90 inhibition was performed using geldanamycin (GA) (blue diamonds). The data for the test compounds (test) are also included (black diamonds). **C.** Chemical structure of the active SEW04784 scaffold. **D.** Chemical structure of SEW04777, an analog that lacked detectable activity in the ASH ATPase assay.

We optimized this assay to monitor the ASH activity in 384-well plate format. Purified Hsp90β and Aha1 were premixed and allowed to reach a binding equilibrium at room temperature. The test compounds were subsequently added and allowed to bind to their target proteins before the addition of ATP and transfer to 37°C to initiate the reaction. After the 3 h incubation, the QR reagent was added and the fluorescence of the white microtiter plates measured. The optimization of the assay revealed a Z’ factor of 0.7, suggesting that the assay was suitable for high throughput screening efforts. We performed our screen using the 16,000 compound Maybridge library, which yielded 131 compounds exhibiting a percent inhibition of the ASH activity exceeding 3 standard deviations of the DMSO treated Hsp90+Aha1 control (**Figure 1B**). A secondary screen to validate these hit compounds reduced this number to 35 target compounds, with our lead compound, <u>SEW</u>04784 (SEW) (**Figure 1C**), being the most potent that did not inhibit the basal Hsp90 ATPase activity (see below). An analog with undetectable activity in the ASH ATPase assay (SEW04777) was used as a reference control (**Figure 1D**).

### Characterization of SEW04784

To characterize SEW, we first performed a titration of the compound to assess its potency in inhibiting the ASH activity. The titration revealed that SEW can completely inhibit this activity at a dose of 5 μM (**Figure 2A**) and exhibits an IC_50_ of 0.3 μM (**Figure 2B**), a value similar to that of the well characterized Hsp90 inhibitor, geldanamycin (GA), which exhibits an IC_50_ value of 0.14 μM (**Figure 2B**). In addition, we observed that while SEW is able to block the ASH (**Figure 2A-C**), it did not affect the basal Hsp90 ATPase activity seen in the absence of Aha1 (**Figure 2A, C**). These data suggest that SEW is a first-in-class compound exhibiting potent inhibitory effect on the co-chaperone stimulated ATPase activity without exerting an inhibitory action on basal Hsp90-allowing the chaperone to function normally in the absence of Aha1-dependent events.

**Figure 2.**
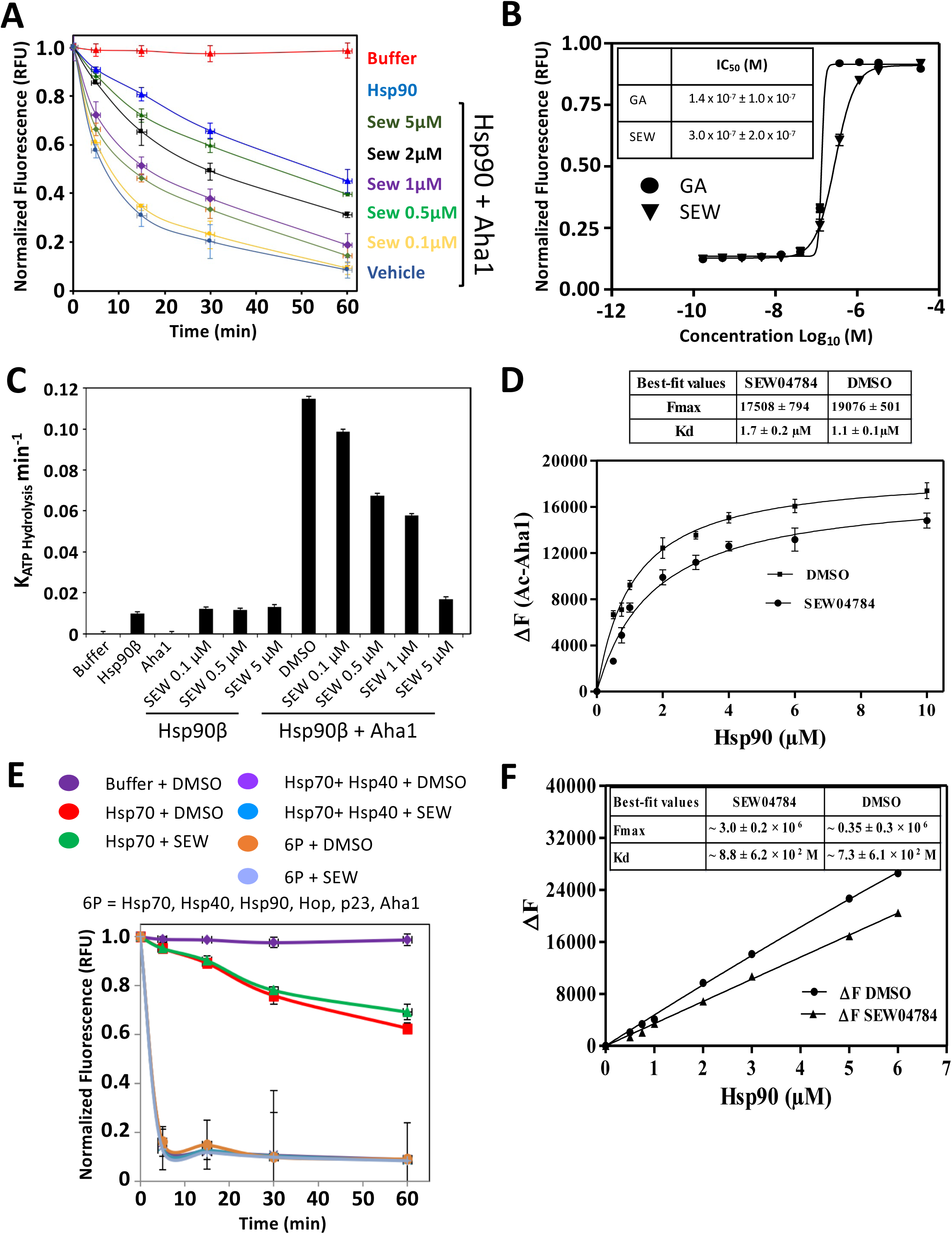
SEW04784 only inhibits the Aha1-stimulated Hsp90 ATPase Activity. **A.** Scatter plot of the normalized fluorescence of the ASH assay using 8 μM Hsp90β and 16 μM Aha1 over time in the presence of DMSO or the indicated concentration of SEW04784 (SEW). **B.** Scatter plot of the normalized fluorescence of the ASH assay using 8 μM Hsp90β and 16 μM Aha1 in the indicated concentrations of GA (circles) and SEW04784 (SEW) (triangles). Inset Table depicts the IC50 values for GA and SEW for the ASH assay. **C.** Bar graph depicting the rate of the ATPase reaction from the ASH assay for the indicated conditions. The amount of Hsp90β and Aha1 used in the assay were 8 μM and 16 μM respectively. **D.** Plot depicting the impact of increasing concentration of Hsp90β on the fluorescence of the acrylodan-labeled full length Aha1 in the presence or absence of SEW04784. The inset Table depicts the Kd and Fmax values for the indicated treatments. **E.** Plot of the normalized fluorescence from QR reagent-based fluorescence quenching monitoring the impact of SEW on the ATPase activity of Hsp70 alone, in the presence of Hsp40 and in the context of the chaperone complex containing Hsp40, Hsp90, Hop, p23 and Aha1 (6P). **F.** Plot of the tryptophan fluorescence of Hsp90 in the presence or absence of SEW04784. The inset Table depicts the Fmax and Kd values for Hsp90β binding to DMSO or SEW.

To corroborate the Aha1-mediated ATPase inhibitory activity of SEW, we addressed the impact of this small molecule on the binding of Hsp90 and Aha1. To monitor the impact of Hsp90 binding on Aha1, we first labeled Aha1 with the molecular probe, acrylodan, whose fluorescence is sensitive to the local environment of labeled cysteine (Cys) residues. In **Figure 2D**, the fluorescence of acrylodan-labeled full length Aha1 (ac-Aha1) increases with increasing concentration of Hsp90 (**Figure 2D**). The addition of SEW to this binding event, significantly reduces the observed increase in ac-Aha1 fluorescence and causes a 50% increase in the KD value (1.7 ± 0.2 μM) for the Hsp90-Aha1 binding reaction relative to that seen in the presence of DMSO (1.1 ± 0.1 μM) (**Figure 2D**). These data suggest that SEW is disrupting the binding of Hsp90 to its regulatory co-chaperone Aha1, consistent with the ability of SEW to block the ASH activity.

In order to assess the specificity of SEW for inhibiting the ATPase activity of Hsp90, we monitored its ability to block ATP hydrolysis of Hsp70, a highly abundant cellular chaperone required for the delivery of client proteins to Hsp90^72^, both alone and in the presence of its ATPase activating co-chaperone Hsp40, as well as in a multi-protein chaperone complex consisting of Hsp70, Hsp40, Hsp90, Hop, Aha1 and p23 (referred to as 6P). The data indicate that SEW does not block the basal ATPase activity of Hsp70 nor did it impact the Hsp40-stimulated ATPase activity of Hsp70 or the ATPase activity of the 6P complex (or multiple combinations thereof (not shown)), where the total ATP hydrolysis is dominated by Hsp70 (**Figure 2E**). These data suggest that SEW is not acting as a general inhibitor of the major core chaperone, Hsp70, responsible for protein folding in the cell.

### SEW04784 binds to the C-terminal domain of Aha1

Since we observed inhibition of the Aha1 dependent Hsp90 ATPase activity in the presence of SEW, we determined if this compound bound directly to Hsp90. We monitored the impact of SEW on the tryptophan (Trp) fluorescence of Hsp90. Here, we observed a linear increase in Trp fluorescence with increasing concentration of Hsp90 in the presence of either DMSO and 25 μM SEW (**Figure 2F**). The presence of SEW had little to no impact on the Trp fluorescence of Hsp90. We estimated a Kd value > 700 M for SEW binding to Hsp90. Taken as a whole, these data suggest that SEW does not bind to Hsp90 based on intrinsic Trp fluorescence response, a conclusion consistent by our observation that SEW does not inhibit the basal Hsp90 ATPase.

In order to address if SEW does bind directly to Aha1, we used transverse relaxation optimized spectroscopy (TROSY) ^1^H-^15^N heteronuclear single quantum correlation (HSQC) NMR. The addition of SEW to full-length Aha1 caused shifts in spectra number of cross peaks in the spectrum, indicating that the compound binds to Aha1 in a localized site (**Figure 3A**). Given that Aha1 is a 2-domain protein that is thought operate as a highly-flexible linked dimer^34,38^, the location of the binding site was further assessed by examining the TROSY-HSQC spectra of the isolated N-terminal (NTD (1-163)) and C-terminal domains (CTD (164-338)) of Aha1 in the presence and absence of SEW. Addition of SEW does not cause any changes in the spectrum of the isolated NTD of Aha1 (**Figure 3B**), but significant chemical shift changes are observed in the CTD of Aha1 (**Figure 3C**). These changes map to the Hsp90 binding interface of the CTD of Aha1 (**Figure 3D**).

**Figure 3.**
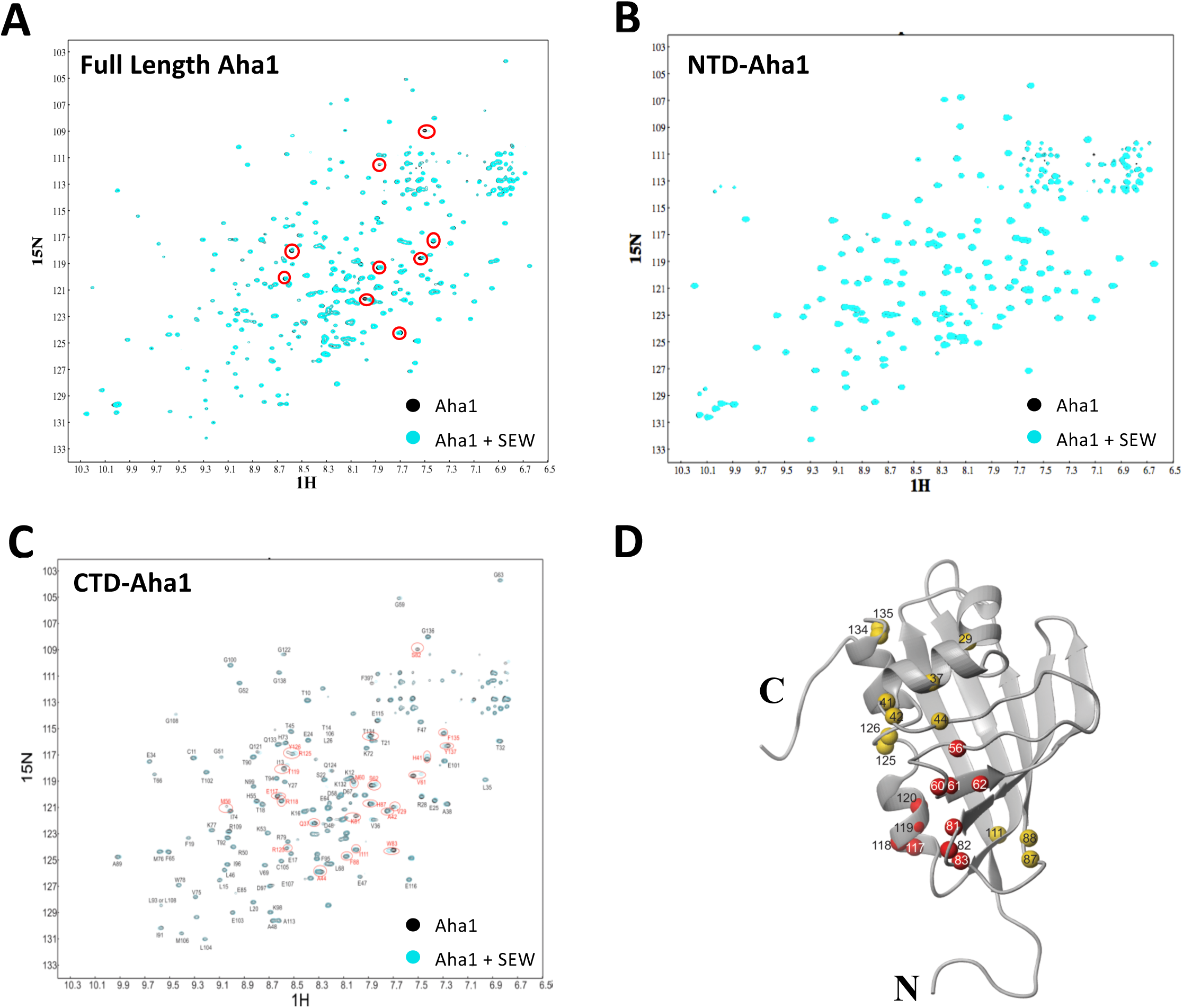
SEW04784 binds to the C-terminal domain of Aha1. **A.** Overlay of 900 MHz ^1^H-^15^N-TROSY-HSQC spectra of full-length Aha1 in the presence of DMSO (black) and an equal amount of a DMSO solution of SEW04784 (blue). Red circles indicate differences in cross peak position between the two spectra. **B.** Overlay of 900 MHz ^1^H-^15^N-TROSY-HSQC spectra of the N-terminal domain (residues 1-163) of Aha1 in the presence of DMSO (black) and an equal amount of a DMSO solution of SEW04784 (blue). **C.** Overlay of 900 MHz ^1^H-^15^N-TROSY-HSQC spectra of the C-terminal domain (residues 164-338) of Aha1 in the presence of DMSO (black) and an equal amount of a DMSO solution of SEW04784 (blue). Assignments are shown and red circles indicate differences in cross peak position between the two spectra. **D.** Ribbon diagram of the C-terminal domain of Aha1 depicting the amino acids exhibiting spectral shifts (red and yellow balls) in response to SEW04784.

In order to confirm that SEW binds to the CTD of Aha1, we monitored the impact of its binding on the fluorescence of acrylodan labeled Aha1. We observed a dose-dependent fluorescence quenching of the acrylodan labeled full length Aha1 (**Figure 4A**) with a Kd of 1.74 μM. We also observed a dose-dependent quenching of the acrylodan fluorescence of Ac-CTD-Aha1 (**Figure 4B**) but not ac-NTD, further supporting the NMR studies showing that the binding of SEW to Aha1 occurs at the CTD of the protein. This conclusion is further supported by our observation that SEW disrupts the binding of the isolated CTD of Aha1 to Hsp90, resulting in a 2.5-fold increase in the Kd for CTD-Aha1/Hsp90 binding (**Figure 4C**).

**Figure 4.**
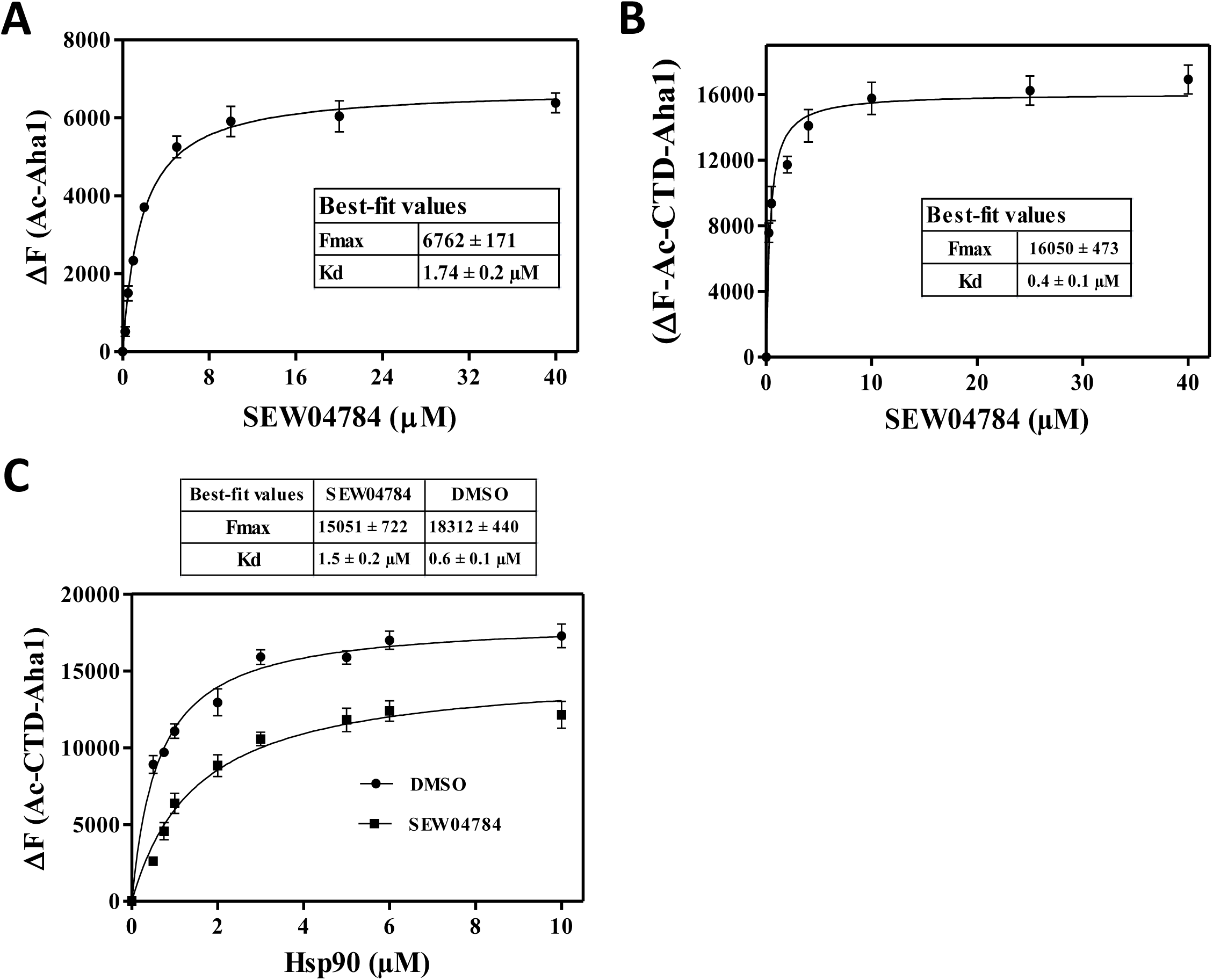
SEW04784 prevents the C-terminal domain of Aha1 from binding to Hsp90. **A.** Plot depicting the impact of increasing concentrations of SEW04784 on the fluorescence of acrylodan-labeled full length Aha1 (ac-Aha1). The inset table depicts the Fmax and Kd values for SEW04784 binding to full length ac-Aha1. **B.** Plot depicting the impact of increasing concentrations of SEW04784 on the fluorescence of the acrylodan-labeled C-terminal domain of ac-Aha1 (164-338). The inset table depicts the Fmax and Kd values for SEW04784 binding to the C-terminal domain of Aha1. **C.** Plot depicting the impact of increasing concentration of Hsp90β on the fluorescence of the acrylodan-labeled C-terminal domain of ac-Aha1 in the presence or absence of SEW04784. The inset Table depicts the Kd and Fmax values for the indicated treatments.

### SEW04784 impedes the re-folding of firefly luciferase

To address whether the ability of SEW to inhibit the ASH activity translates to an inhibition of the chaperoning properties of Hsp90, we monitored the effect of SEW on the refolding of firefly luciferase (FLuc) *in vitro*. Thermal denaturation of FLuc destroys its ability to oxygenate luciferin, which translates to the loss of luminescence originating from oxyluciferin. This process can be partially reversed by the addition of the 6P chaperone complex (**Figure 5A**). The refolding of FLuc is dependent on the presence of the Hsp70 chaperone, involved in client delivery to Hsp90, as evidenced by the ability of the Hsp70-inhibitor, JG98^73^, to completely block the refolding of FLuc. The refolding potential of the 6P chaperone machinery is also dependent on the presence of Hsp90 during the thermal denaturation of FLuc as evidence by the impact of geldanamycin (GA), an Hsp90 ATPase inhibitor (**Figure 5B**). Inhibition of Hsp90 with GA reduces the folding capacity to 75% of the control lacking GA. We observed that SEW causes similar reduction in the refolding capacity of the 6P system, suggesting that management of Hsp90 sensitive refolding of Fluc is dependent on the SEW sensitive activity of Aha1.

**Figure 5.**
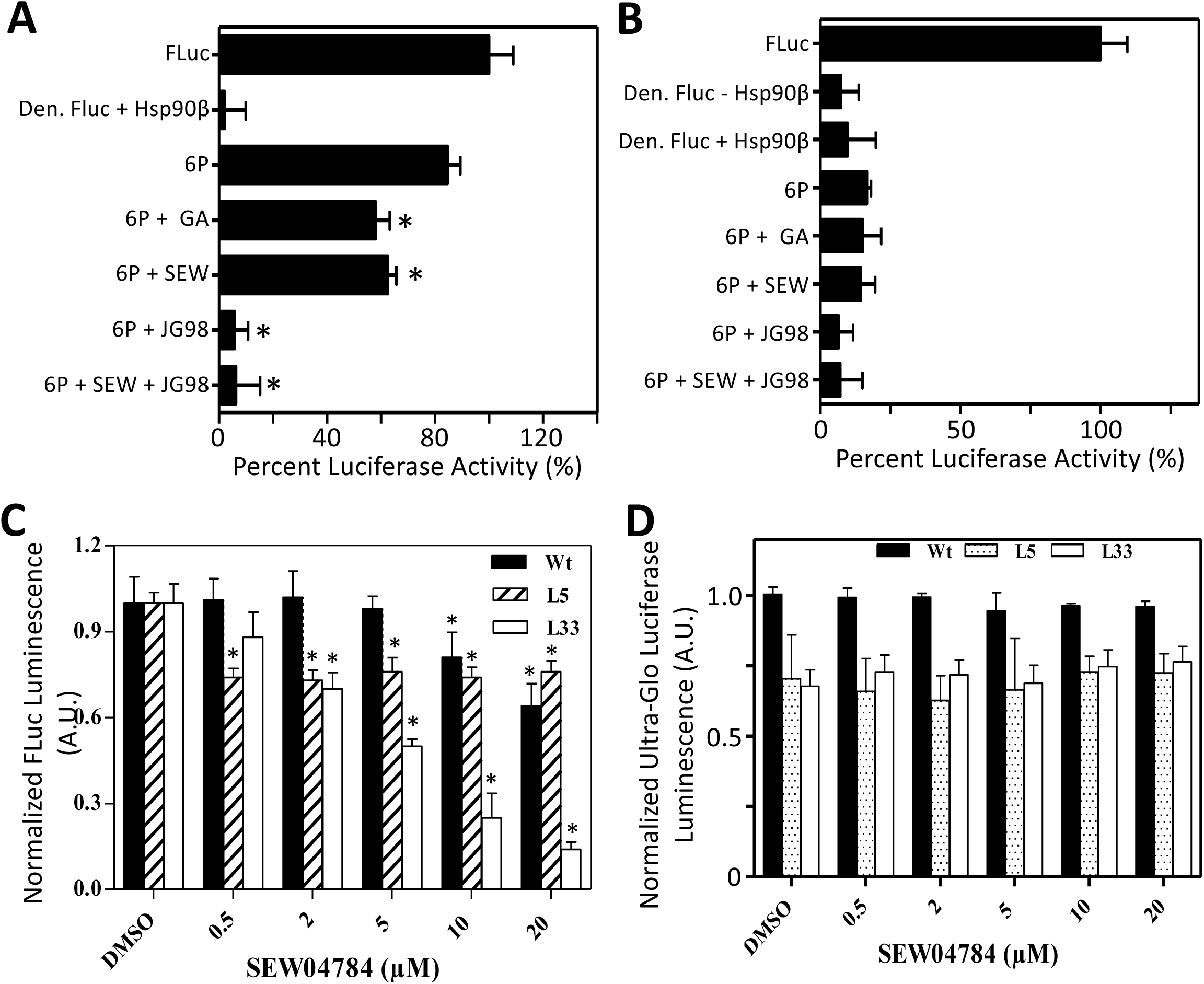
SEW04784 inhibits the Hsp90-dependent folding of Firefly Luciferase. **A-B.** Bar graph depicting the normalized activity of Firefly Luciferase (FLuc) in response to *in vitro* refolding of recombinant, purified FLuc in the presence of the indicated chaperone proteins following heat denaturation in the presence (**A**) or absence (**B**) of Hsp90β. The 6P protein complex is composed of Hsp90, Hsp70, Hop, p23, Aha1 & Hsp40. JG98 and GA are well characterized Hsp70 and Hsp90 inhibitors, respectively. The data is shown as percent (%)Fluc activity relative to the activity of FLuc prior to thermal denaturation at 45ºC for 8 min. **C.** Bar graph depicting the normalized FLuc luminescence of transiently transfected HeLa cells with FLuc variants (WT, L5 (R188Q) and L33 (R188Q; R261Q) in the presence of the indicated concentration of SEW04784. **D.** Cellular toxicity assay depicting the impact of the indicated concentration of SEW04784 on the cellular ATP levels of HeLa cells transfected with the indicated FLuc variant as measured by the CellTiter-Glo luminescence assay. The data is normalized to the luminescent signal obtained in WT-transfected HeLa cells treated with vehicle (DMSO).

In order to capture the impact of SEW *in vivo*, we assessed its impact on cells stably expressing variants of the FLuc. In addition to the WT-FLuc, we studied 2 variants, L5-FLuc (R188Q) and L33-FLuc (R188Q/R261Q), that contain mutations that disrupt polar contacts in the 3-dimensional (3D) structure of the protein, thereby affecting their stability^74^. SEW caused a dose-dependent decrease in WT-FLuc activity above a dose of 5 μM (**Figure 5C**). This effect was more pronounced for the folding sensitive L33-FLuc variant, where we again observed a dose-dependent destabilization of the protein at a dose as low as 0.5 μM (**Figure 5C**). The L5 variant was also sensitive to SEW at a dose of 0.5 μM but retained significantly more activity than that seen with the L33 variant at increasing doses (**Figure 5C**). This SEW-dependent loss of folding of FLuc variants was not due to cytotoxicity since we did not observe any toxicity at the doses tested using the CellTiter Glo assay (**Figure 5D**). While the expression of the mutant FLuc variants cause a 25% reduction in cell viability relative to that seen in cells expressing the WT-FLuc, the addition of SEW to these cells did not further promote cytotoxicity (**Figure 5D**). Therefore, it appears that SEW is capable of inhibiting the chaperoning activity of Hsp90 in the complex environment of the cell.

### SEW04784 inhibits the activity of steroid hormone receptors (SHR)

In order to determine the impact of SEW on biologically relevant Hsp90 client proteins, we turned our attention to the steroid hormone receptor family (SHR). We assessed the ability of SEW to inhibit the transcriptional activity of two SHRs, namely the glucocorticoid receptor (GR) and androgen receptor (AR). The binding of cognate ligands induces nuclear translocation of SHR to induce transcription of genes responsive to a given receptor/ligand pair. We utilized a stably integrated reporter construct composed of the FLuc gene under the control of a GR-(**Figure 6A**) and AR- (**Figure 6B**) responsive promoter in MDA-kb2 cells. The addition of the cognate ligand activates the expression of the FLuc gene, whose luminescence can be monitored after the addition of luciferin.

**Figure 6.**
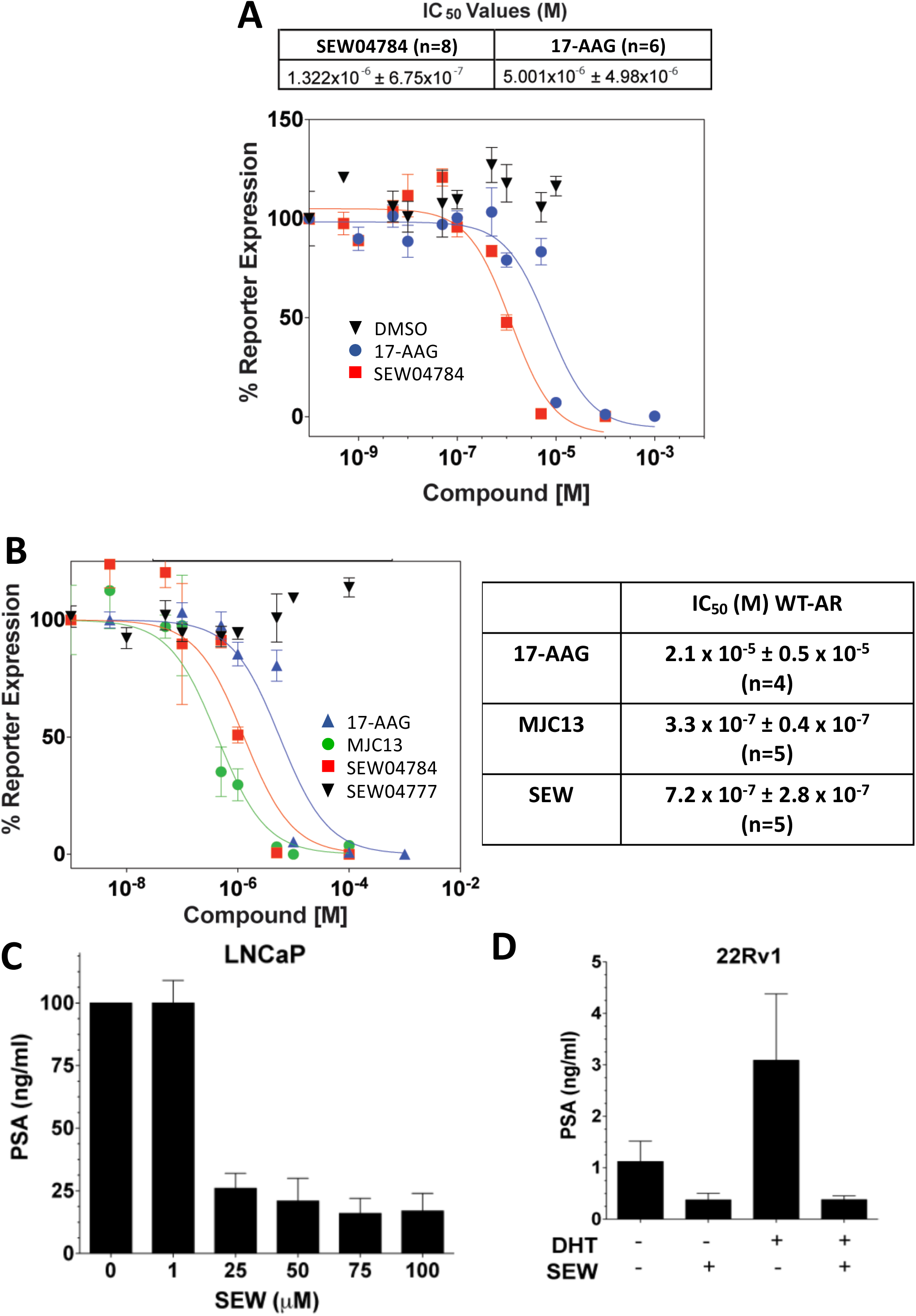
SEW04784 inhibits the transcriptional activity of steroid hormone receptors. **A.** Sigmoidal inhibition curves showing the impact of SEW04784 (red squares) and 17-AAG (blue circles) on the transcriptional activity of the glucocorticoid-stimulated glucocorticoid receptor (GR) in cells expressing a FLuc reporter construct under the control of a GR-responsive promoter. The table depicts the IC50 values for SEW04784 and 17-AAG. **B.** Sigmoidal inhibition curves showing the impact of the Hsp90 inhibitors, SEW04784 (red squares), 17-AAG (blue triangles) and MJC13 (green circles) as well as the SEW analog, SEW04777 (black triangles) on the transcriptional activity of the dihydrotestosterone (DHT)-stimulated WT-androgen receptor (AR) in cells expressing a FLuc reporter construct under the control of a AR-responsive promoter. Table depicting the IC_50_ values for SEW04784, 17-AAG and MJC13 from the AR experiment. **C**. Bar graph showing the impact of the indicated concentrations of SEW04784 on the endogenous activity of the T877A-AR variant in LNCaP cells, a prostate cancer associated AR variant, in the presence of 50 nM DHT. The activity of the T877A variant was assessed by measuring the level of secretion of the prostate specific antigen (PSA), an AR-responsive gene found in prostate cells. **D.** Bar graph showing the impact of 100 µM SEW04784 on the endogenous activity of the AR variants in 22Rv1 cells. These cells express both a WT-AR allele as well as an AR variant lacking its ligand binding domain (ΔLBD)-a constitutively active AR variant that does not require DHT for activation of its transcriptional activity. The activity of these 2 variants were separated by monitoring the secretion levels of PSA in the absence and presence of 1 nM DHT.

We observed that SEW inhibited the GR-induced luciferase expression with an IC_50_ of 1.3 μM, a value similar to that seen for the Hsp90 inhibitor 17-AAG (5.0 μM) (**Figure 6A**). In agreement with this observation we also noted that SEW was able to inhibit the WT-AR-induced expression of Fluc with an IC_50_ value of 0.7 μM^75^, a value similar to that seen for the FKBP52-specific AR inhibitor MJC13 (0.3 μM) (**Figure 6B**), and well below that for the Hsp90 inhibitor 17-AAG (21 μM) (**Figure 6C**). Additionally, the SEW analog SEW04777 with undetectable activity (**Figure 1D**), did not impact the AR-dependent expression of the Fluc reporter (**Figure 6B**).

Mutations in the AR can cause significant changes in the behavior of this SHR leading to transformation of affected cells, culminating in prostate cancer (PCa). The T877A mutation of the AR, results in a receptor that can be activated by additional steroids, including estrogen and anti-androgens, allowing prostate cells that express this variant to continue to grow in the face of androgen deprivation therapy (ADT) and chemical castration by responding to other steroids or anti-androgens^76-81^.

To assess the therapeutic potential of SEW in treating PCa-associated AR variants, we monitored the impact of SEW on LNCaP cells, a prostate cancer cell line that is homozygous for the T877A-AR variant. Here, we observed a dose-dependent inhibition of expression and secretion of the prostate specific antigen (PSA) (**Figure 6C**), an AR-responsive gene used as a biomarker for the onset and progression of PCa. In addition to missense mutations, deletion of the ligand binding domain (ΔLBD) of the AR leads to constitutively active variants of the AR responsible for aggressive forms of PCa. These mutations are insensitive to ADT and are currently without treatment. To address if SEW-mediated inhibition of Hsp90 would have any impact on ΔLBD variants, we monitored its impact on the heterogeneous 22Rv1 prostate cancer cell line that express both a dihydrotestosterone (DHT) responsive variant and ΔLBD-AR variant. SEW was able to inhibit both the constitutive and DHT-responsive secretion of PSA (**Figure 6D**) at a dose of 20 μM, suggesting that it could provide significant therapeutic benefit in the treatment of both androgen-dependent and castration resistant PCa (CRPC).

### SEW04784 can reduce the expression of phospho-tau

Tau is a microtubule associated protein (MAP) that promotes and stabilizes the formation of axonal microtubules^82,83^. Mutations or hyperphosphorylation of tau (p-tau) leads to altered microtubule binding culminating in the loss of axonal transport^84,85^ and tau aggregation^86^. This latter effect is a hallmark of a series of neurodegenerative diseases termed tauopathies, the most prominent of which is Alzheimer’s disease (AD), where p-tau aggregates can be found in neurofibrillary tangles, senile plaques and cellular processes^87-91^. p-tau aggregates have been shown to be sensitive to the protective action of the heat shock protein molecular chaperone system^91^. Recent evidence has demonstrated that small molecules inhibitors of both Hsp70 and Hsp90, or modulation of the expression of regulatory co-chaperones, can mediate the clearance of p-tau in cells^87-97^.

In order to address the ability of SEW to specifically target toxic p-tau species, we utilized a HeLa cell line stably expressing WT-tau(4R0N). Here, we observed that SEW selectively cleared p-tau, specifically pS396/pS404, in a dose dependent manner, where a complete elimination of this toxic species could be observed at a dose of 30 μM (**Figure 7A**). The analog of SEW that had undetectable activity in the ASH ATPase assay, SEW04777 (**Figure 1D**), did not affect the levels of p-tau, suggesting that the observed effects of SEW are related to its ASH ATPase inhibitory activity. Interestingly, SEW had a less pronounced effect on the clearance of total tau (**Figure 7A**), a result consistent with previous observations showing that Hsp90 inhibitors can selectively clear specific p-tau species^93^. The treatment of HeLa-tau(4R0N) cells with a dose of 10 μM SEW demonstrated that the effects could be seen as early as 1 h post-treatment and continue to promote the clearance of p-tau (pS396/S404) throughout the 24 h dosing regimen (**Figure 7B**).

**Figure 7.**
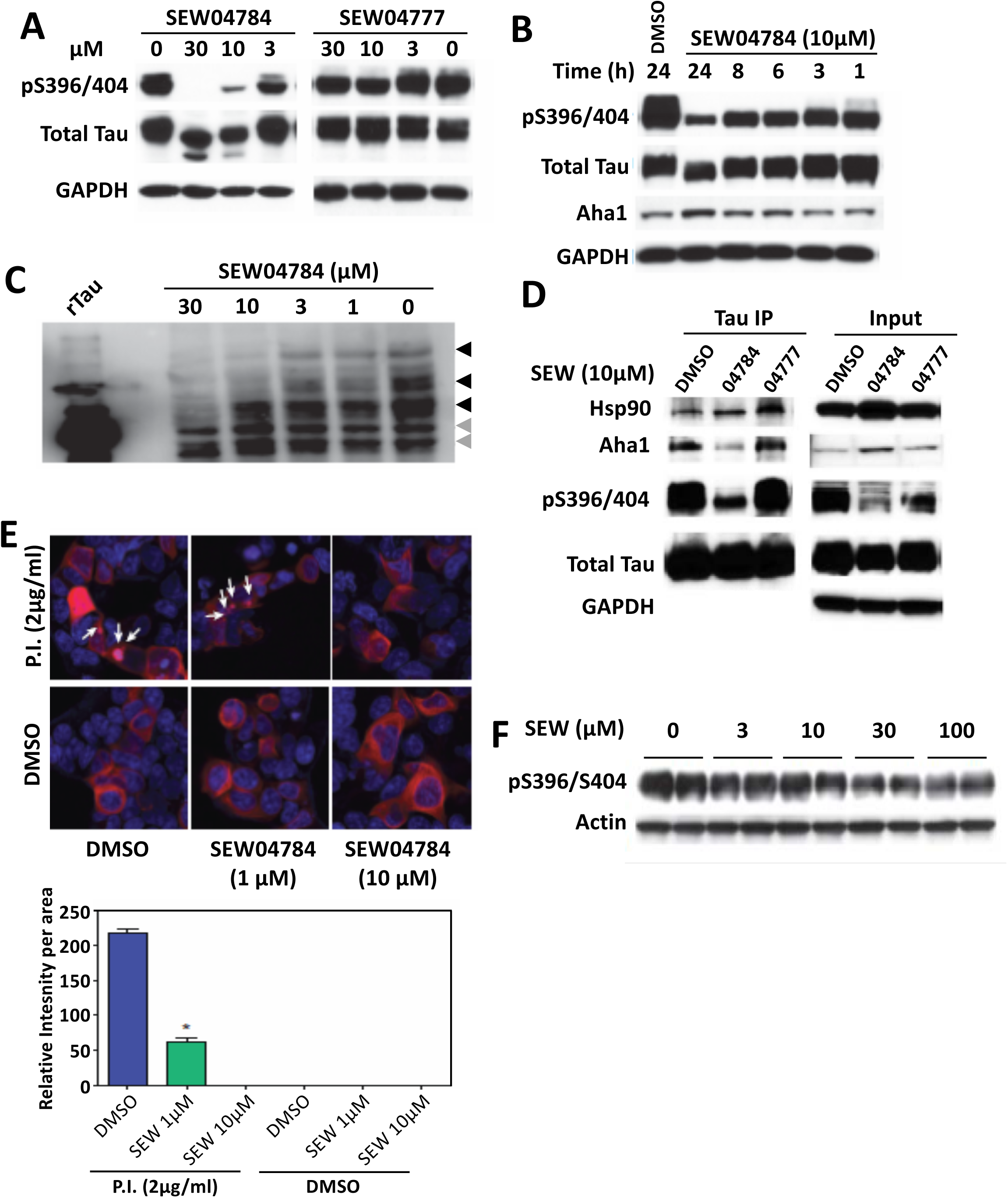
SEW04784 clears toxic phosphorylated tau species. **A.** Western blot analysis of lysates prepared from HeLa cells stably expressing tau-4R0N following treatment with the indicated concentration of SEW04784 or SEW04777. The immunoblot was probed with antibodies specific for total tau and phosphorylated tau (pS396/404) (PHF) as well as for GAPDH. **B.** Western blot analysis of lysates prepared from HeLa cells stably expressing tau-4R0N following treatment with 10 μM SEW04784 for the indicated time. The immunoblot was probed with antibodies that recognize total tau, phosphorylated Tau (S396/404) (PHF), Aha1 and GAPDH. **C.** Immunoblot analysis of lysates prepared from HeLa cells stably expressing tau-4R0N following treatment with the indicated concentration of SEW04784 for 24h. The lysates were separated on a Phos-Tag PAGE and the immunoblot analyzed using an antibody that detects total tau. Recombinant tau (rTau) was included as loading control to distinguish between non-phosphorylated tau (grey arrowheads) and phosphorylated tau (black arrowheads). **D.** Immunoblot analysis of tau immunoprecipitates (left) from HeLa cells lysates expressing tau-4R0N treated with SEW04784 (04784), SEW04777 (04777) or vehicle (DMSO). The input lysates (right) were also analyzed. The immunoblots were analyzed with antibodies that detect Hsp90, Aha1, phosphorylated tau (pS396/404) (PHF), total tau and GAPDH. **E.** Immunofluorescence (upper panel) of HEK293 cells transiently transfected with RFP-tau-4R0N and treated with 2 μg/ml protease inhibitor (P.I.) or vehicle (DMSO) in the presence or absence of the indicated concentration of SEW04784. The white arrowheads indicate the location of RFP positive tau aggregates. The quantitative analysis (lower panel) provides the relative intensity per area for aggregated RFP-tau in the immunofluorescence images on the left panel for the indicated treatments. **F.** Western blot analysis of lysates prepared from cultured brain slices prepared from transgenic mice expressing the P301L-tau variant. The immunoblot was analyzed with antibodies that detect phosphorylated tau (pS396/S404) and actin^125^.

To confirm the observation that SEW is promoting the preferential clearance of p-tau species, we employed the Phos-tag PAGE approach^98,99^, which allows the separation of phosphoproteins on acrylamide based the degree of phosphorylation. **Figure 7C** shows a dose-dependent clearance of the p-tau species (black arrows) relative to the non-phosphorylated species (grey arrows) with complete clearance observed at 30 μM (**Figure 7C**). To gain further insight in the mechanism of action of SEW, we performed an immunoprecipitation of tau from HeLa-tau(4R0N) cells to assess if SEW was impacting the binding of Hsp90 and/or Aha1. We observed that the active SEW analog (SEW4784) altered the recruitment of Aha1 to tau complexes without affecting the binding of Hsp90 (**Figure 7D**), although we observed a slightly increased binding of Hsp90 to tau. Conversely, the SEW04777 analog lacking measurable ASH ATPase inhibitory activity, did not affect the recruitment of Hsp90 or Aha1 (**Figure 7D**).

Given that tau phosphorylation is linked to aggregation, we used a well-characterized cell model expressing fluorescently-tagged tau, which allows for monitoring the aggregation in response to proteasome inhibition^100^. The treatment of HEK cells, transiently transfected with RFP-tagged WT-4R0N tau, with proteasome inhibitor (P.I.) leads to the formation of RFP-positive aggresomal clusters, which are not seen in vehicle treated cells (**Figure 7E**). The co-administration of SEW at doses as low as 1 μM significantly abrogates the aggregation of tau (**Figure 7E**), an effect which can be completely blocked at high dose of SEW (**Figure 7E**).

To provide secondary validation in another physiologically relevant model, we tested the effect of SEW on p-tau levels from cultured brain slices from a 4 month old transgenic mouse (rTg4510), which expresses the P301L variant of tau, a mutation shown to disrupt microtubule binding^84,101^ and thereby increases the aggregation propensity of tau. We observed a dose-dependent reduction in the levels of p-tau (pS396/S404) in mouse brain tissue (**Figure 7F**), suggesting that SEW can provide protection to these toxic tau species.

## Discussion

Herein, we describe the discovery and characterization of a novel inhibitor of the Aha1-stimulated Hsp90 (ASH) ATPase activity, SEW04784 (SEW). SEW binds directly to the Hsp90 co-chaperone Aha1 protein to inhibit its ATPase stimulating activity. Inhibition of co-chaperone activity alters the folding behavior of client proteins such as FLuc, SHRs and p-tau aggregates, highlighting the potential versatile therapeutic value of this small molecule inhibitor for Hsp90 sensitive client specific folding.

### Regulation of SHR function

The chaperoning activity of Hsp90 was first characterized using the well characterized family of SHRs^102,103^ including AR involved in prostate cancer. PCa is characterized by dysregulated androgen and AR-dependent growth of prostate cells, often associated with AR mutations. The standard of care for this disease includes androgen depletion therapy (ADT), which includes chemical or physical castration, to remove the source of androgens and abrogate the growth of prostatic tissue. ADT eventually fails resulting in a castration resistant state due to AR overexpression, mutations conferring ADT resistance, enhanced co-regulator expression and/or activity and intratumoral production of androgen. While the treatment with abiraterone, a CYP17A1 inhibitor to block the intratumoral synthesis of androgen^104^, and enzalutamide, an AR-LBD antagonist^105^, are designed to slow the growth of CRPC, many patients are refractory to these treatments or quickly develop resistance^106^, possibly due to development of AR truncation mutants lacking the LBD, which exhibit constitutive activity. Therefore, the development of therapeutic options which can inhibit androgen responsive AR variants in the early stages of the disease as well as androgen-resistant AR variants associated with CRPC are critical for extending the lifespan of affected individuals. A further level of complication arises from the observations that in many cases of CRPC with AR truncation variants, there is a concomitant increased expression of the GR which has the capacity to modulate the expression of many AR responsive genes in prostate cells^107-110^, thus providing a way for these transformed cells to escape growth arrests mediated by AR inhibitory therapies. Therefore, identifying therapeutic targets that are critical for the functionality of SHR and developing small molecules that can inhibit these growth promoting factors or their associated signaling pathways are of high importance as shown herein for Aha1 inhibitor scaffold SEW that inhibits both AR and GR activity *in vivo* by its interaction with Aha1.

While ΔLBD-AR variants have been shown to be refractory to Hsp90 inhibition^111,112^, due to the fact that the Hsp90 binds to the AR via the LBD, we demonstrated that SEW is also able to inhibit both the constitutive and DHT-mediated expression of PSA in 22Rv1 cells that express both a WT-AR allele and the CRPC-associated AR-V7 ΔLBD variant, suggesting that an Hsp90-independent activity of Aha1, which is also sensitive to SEW04784, might contribute to the mechanism of inhibition of this subclass of PCa-associated AR variants. In addition, recent evidence suggests that the Hsp40-Hsp70 cycle is intimately involved in management of AR and a chalcone scaffold series that blocks AR function in promoting proliferation of numerous prostate cancer models operate through the Hsp70 co-chaperone Hsp40^71^. These results illustrate the importance of the entire Hsp70-Hsp90 chaperone/co-chaperone cycle in client delivery and management of misfolding as drivers of prostate cancer progression.

### Role of Aha1 in tauopathy

Hsp90 is critical for the biogenesis and functional cycling of numerous clients proteins in addition to SHR (https://www.picard.ch/Hsp90Int/index.php)^5,113,114^, with microtubule associated protein, tau, being one of the best characterized^87-91,115^. Small molecule inhibitors of Hsp90 have emerged as potential therapeutic options for the treatment of tauopathies^87,89,90,116^. While elimination of toxic aggregates has been demonstrated at the bench, translation to the bedside has been disappointing, largely due to the expected associated cytotoxicity given the central role of Hsp90 in proteostasis.

To circumvent the toxic effects of Hsp90 ATPase inhibitors, recent efforts have turned to targeting the Hsp90 co-chaperone machinery. The molecular silencing of most Hsp90 regulatory co-chaperones impedes the ability of 90inh to mediate the clearance of toxic p-tau species ^93^, suggesting that it is the Hsp90 chaperone/co-chaperone complex that facilitates clearance and not Hsp90 in isolation. Indeed, Aha1 is critical for the Hsp90-mediated aggregation of p-tau species and that inhibiting its Hsp90 binding with small molecules targeting Hsp90 to prevent binding to Aha1^17,20^, or those that are presumed to interact directly with Aha1 cannot only prevent the formation of de novo aggregates but also promote the clearance of p-tau^18^. Consistent with these observations, SEW04784 binds the C-terminal domain of Aha1 and prevents the aggregation of toxic tau species and selectively promote the clearance of phosphorylated, aggregation prone species (S396/S404). Taken together, these data suggest that the normal Aha1-dependent cycle of Hsp90, serves to protect phosphorylated soluble tau species, leading to their subsequent aggregation and eventual formation of neurofibrillary tangles characteristic of tauopathies. Alterations in the ability of Hsp90 to bind ATP, as seen with 90inh such as GA and 17-AAG, to adopt a stabilized ATP-bound state, as seen with p23 silencing^93^, or to hydrolyze ATP, as seen with a potential selective inhibitor of Aha1 binding directly to Aha1 as shown herein, lead to the clearance of Hsp90-Aha1 bound tau species^92^.Inhibiting Aha1 directly has the benefit of not altering the basal chaperoning function of Hsp90, thereby mitigating the cytotoxic issues seen with complete Hsp90 inhibition^58,59^. Thus, a novel, specific and potent Aha1 inhibitor presents a potential therapeutic opportunity for the treatment of neurodegenerative diseases upon further SAR of the parent SEW scaffold (B. Tait and J.E. Gestwicki, not shown).

While the cellular function of Aha1 is often exclusively considered in the context of its co-chaperoning activity as an ATPase stimulating co-factor for Hsp90, the majority of Aha1 exists in an Hsp90-free state in human cells^25^. While this mutual exclusion observation could be explained by the co-chaperone activity of Aha1, which shifts the binding equilibrium of the Hsp90/Aha1 interaction to the non-bound state, Aha1 was able to prevent the aggregation of both FLuc and Rhodanese *in vitro* and *in vivo* in the absence of Hsp90, suggesting that it possesses direct chaperoning activity. While the ability of Aha1 to bind to its clients has been shown to require the first 22 amino acids located at the NTD of Aha1^17,39^, which is unaffected by SEW, the ATPase stimulating activity *in vitro* required the full-length protein. These data suggest that while SEW04784 is unlikely to disrupt the ability of Aha1 to bind to its client proteins, it potentially could also impact the chaperoning activity of Aha1, thereby promoting the clearance of bound clients, such as p-tau and the AR, through ubiquitination, as shown tau in response to co-incubation with IU1-47^117^ or for Aha1-bound, heat denatured FLuc in the absence of chaperone-mediated refolding^25^. Therefore, the ability of SEW to mediate a reduction in tau aggregation and/or block the activity of AR missense variants in prostate cancer could be the concerted action of both inhibiting the ASH activity and the chaperone activity of Aha1. Its impact on truncated AR variants may be explained by its latter function when AR is lacking the Hsp90 interaction site.

Our results highlight the potential benefits of Aha1 selective inhibitors in managing human misfolding disease. Such compounds allow the cell to maintain the basal activity of Hsp90, mitigating the cytotoxic effects reported with the use of direct Hsp90 binding inhibitors. We have recently suggested by applying variation spatial profiling (VSP) to analyze the impact of variation in individual based on the world-wide population, that the cellular environment is critical in setting spatial covariance (SCV) tolerance by the cell in response to the environment^16^. These results lead us to define Aha1 specific regulators as SCV managers that modify the impact of inherited and environment induced misfolding stress on the fold. SCV^16^, as an ancient^7^ proteostasis sensitive platform for management human misfolding disease^8,12^, provides a new general paradigm for understanding disease in the more global context of SCV-setpoints that may support sequence-to-function-to-structure relationships in aging, health and disease^1,6-12,16^.

## Materials and Methods

### Expression and purification of human Hsp90β

An overnight culture of BL-21 (DE3) *E.* coli carrying the pET14b plasmid containing the human Hsp90β (pET-90β) cDNA was diluted into 2 liter of Luria Broth (LB) medium containing carbenicillin (100 g/ml) and grown at 37°C to an optical density (OD) of 0.8 (∼5 h). The *E. coli* cells harboring pET-90β plasmid were induced by 1 mM isopropyl β-D-thiogalactoside (IPTG) at 20°C for 16 h. Pelleted cells were resuspended in 40 ml of immobilized metal affinity chromatography (IMAC) buffer A (20 mM NaH2PO4 pH 8,500 mM NaCl, 1 mM MgOAc, and 5 mM β-mercaptoethanol (β-ME)) supplemented with a EDTA-free protease inhibitor tablet (Roche Diagnostics). The cells were then lysed on ice by sonication at 40% power for 3x for 20 s with a 30s recovery between each sonication burst. The lysate was centrifuged at 25 000xg for 50 min in a type 70 Ti rotor (Beckman Coulter), and the supernatant was fractionated over a 5 ml Ni^2+^-charged HiTrap Chelating HP column (GE Healthcare) using an AKTA FPLC with Frac-950 fraction collector (GE Healthcare). Gradient fractionation was carried out with IMAC buffer B (IMAC A with 1 M imidazole). Hsp90β containing fractions were pooled, concentrated using an Amicon Ultra-15 Centrifugal Filter Unit with Ultracel-50 membrane, MWCO: 50,000 (Millipore), and further gradient fractionated on an 8-ml Mono-Q HR 10/10 column, (GE Healthcare) using Mono-Q buffer A (20 mM Tris, pH 7.5, and 1 mM DTT), and Mono Q buffer B (20 mM Tris, pH 7.5, 1 M NaCl, and 1 mM DTT). Hsp90β containing fractions were collected and concentrated to a minimal volume (0.5 ml) using an Amicon Ultra-15. The concentrated sample was passed through gel filtration chromatography (GFC) using Hi-load™ 16/60 Superdex 200 prep grade (GE Healthcare) in 40 mM HEPES/KOH, pH 7.5, 300 mM KCl, 1 mM EDTA, and 1 mM DTT. Final purity was ≥98% based on SDS-PAGE.

### Expression and purification of human Aha1

An overnight culture of BL-21 (DE3) *E.* coli carrying the pET14b plasmid containing the human Aha1 (pET-Aha1) cDNA was diluted into 2 liter LB medium containing carbenicillin (100 µg/ml) and grown at 37°C to an OD of 0.8 (∼5h). The cells were induced by 0.5 mM IPTG at 30°C for 5 h. After induction, the cells were pelleted and stored at −80°C. Pelleted cells were resuspended in 40 ml of immobilized metal affinity chromatography (IMAC) buffer A (50 mM Tris-HCl pH 8, 500 mM KCl, and 5 mM β-ME) supplemented with a protease inhibitor tablet. Cell lysis and HiTrap Chelating HP column and Mono-Q column purification were performed as described above. Aha1 containing fractions were pooled, concentrated, and further fractionated by high performance gel filtration chromatography using a Superdex™75 HR 10/30 (Amersham Biosciences) in 40 mM HEPES, pH 7.5, 50 mM KCl, 2 mM MgCl_2_ and 1 mM DTT. Final purity was ≥98% based on SDS-PAGE. Purified Hsp90β and Aha1 were aliquoted and snap-frozen in liquid Nitrogen prior to storage at - 80°C.

### Expression and purification of human Hsp70, Hsp40, Hop, Bag and p23

Hsp70 was purified as described with minor modifications^118^. Bag1 was purified as described with minor modifications^119^. HOP was purified as described with minor modifications^120^. p23 was purified as described with minor modifications^121^.

### Quinaldine Red Assay

Based on pioneering efforts of Gestwicki ^71,122^, we developed a modified quinaldine red (QR) based HTS assay for screening of compounds that specifically inhibit Aha1. Quinaldine red (QR) reagent was made by mixing 1 g ammonium molybdate, dissolved in 14 ml of 4N HCl, 43 mg QR dye dissolved in 40 ml of 1N HCl and solutions, 1.2ml of 1% poly vinyl alcohol (PVA) and completing the volume to 100 ml with ddH_2_0 water. To initiate the QR assay, 2.3 µM Hsp90 and 14 µM Aha1 were added to the wells of a 384-well white microtiter plate (Greiner Bio-one) in reaction buffer (25 mM Hepes pH 7.5, 10 mM MgCl_2_, and 1 mM DTT). The plate was subsequently centrifuged 250xg for 1 min and the proteins allowed to incubate at room temperature for 30 min. Subsequently, 50 nl of the test compounds from the Maybridge library (or DMSO vehicle) were pinned into the appropriate wells, the plate centrifuged at 250xg for 1 min and incubated at room temperature for 30 min. The reaction was initiated by adding 14mM final concentration of ATP, the plate centrifuged at 250xg for 1 min and incubated at 37oC for 3 h in a humidified environment. Following the 3 h reaction, plates were transferred to room temperature for 10 min and then 15 µl of QR reagent was added to each well of the plate using a Multidrop Dispenser. The ability of the QR reagent to quench the inherent fluorescence (Ex/Em 430nm/530nm) of the white microtiter plates was measured using an EnVision Multilabel Fluorescence Reader (PerkinElmer). The signal to background (S:B) ratio and Z’ factor were determined in triplicates in 384-well format. The S:B ratio was calculated using the equation: S:B = median of high control/median of low control. The low controls are those wells that have Hsp90, Aha1, ATP, and DMSO in reaction buffer, which will generate the highest amount of Pi and therefore provide the highest QR reagent-mediated quenching of fluorescence. The high controls are wells that have Aha1, ATP, and DMSO in buffer in the absence or presence of compound since they will generate the least amount of Pi and therefore provide little quenching of the fluorescence of the microtiter plates. Z’ was calculated based on the equation: Z’ = 1 – 3(*σ*_H_ + *σ*_L_)/(μ_H_ - μ_L_), where *σ* and μ represent the standard error and mean for the High (H) and Low (L) controls. The percent inhibition was defined as 100 – (High Control – Test condition)/(High Control – Low Control). Inhibitors of the Aha1-stimulated Hsp90 ATPase were defined as those with percentage inhibition greater than 3 X standard deviation from the low control. To validate the QR assay, we also established conditions to measure human Hsp90 ATPase activity using the Promega ADP Glo Max™ Kinase Kit according to the manufacturer’s directions with identical results (data not shown).

### Measurement of effect of SEW04784 on the ability of Aha1 to inhibit the ATPase activity of Hsp90

Hsp90 (8 µM) was added to the wells of a 384-well plate containing reaction buffer (25 mM Hepes pH 7.5, 10 mM MgCl_2_, and 1 mM DTT) and incubated on ice for 30 min in the absence and presence of Aha1 (16 µM). Following the incubation period, different concentrations of SEW04784 (0, 0.1, 0.5, 1, and 5 µM) were added to the reaction mixture and the reaction allowed to continue for an additional 30 min on ice. Subsequently, 2 mM ATP was added to the reaction mixture and the plate immediately transferred to 37ºC to initiate ATPase activity. To assess the rate of ATP hydrolysis, 7 µl aliquots were withdrawn at 0, 3, 6, 15, 30, and 60 min and transferred to the wells of a white 384 well microtiter plate containing 3 µl of EDTA (100 mM) on ice. After completion of the time course, 15 µl of the QR reagent was added to each well followed immediately by 2.5 µl of 35% sodium citrate and mixed. Kinetics of Aha1 stimulated Hsp90 ATP hydrolysis was measured using a BioTek Synergy Mx Monochromator based multi-mode plate reader.

### Labeling of Aha1 with acrylodan

The cysteine residues of human Aha1 were chemically modified by a thiol reactive acrylodan (6-Acryloyl-2-Dimethylaminonaphthalene) molecular probe ^123^. For acrylodan labeling, Aha1 was incubated in 40 mM Hepes, pH 8.0, 50 mM KCl, 5 mM MgCl_2_ with 5-fold molar excess of acrylodan at 4 °C for 20h. The reaction mixture was spun at 40000xg for 30 min at 4°C to remove any aggregate formed during reactions. The unbound acrylodan were removed by dialysis against 20 mM Hepes buffer, pH 7.5, 10 mM KCl and 1 mM DTT at 4°C for 24 h, with buffer changes every 8 h. The concentration of Aha1 was measured at 280nm, and the concentration of Aha1-bound acrylodan (Acrylodan-Aha1) was determined by using an extinction coefficient of 20,000 M-1 cm-1 at 392 nm. Incorporation stoichiometry was calculated by dividing acrylodan-Aha1 concentration by the Aha1 concentration. The labeling efficiency of acrylodan to Aha1 was found to be 0.8 ± 0.15.

### SEW04784 does not significantly perturb the dissociation constant of Hsp90β and Aha1 complex

Acrylodan-Aha1 (1 µM) was incubated, in the absence and presence of Hsp90 (0 to 6 µM) for 30 min over ice in reaction buffer (20 mM Hepes, pH 7.5, 10 MgCl_2_, 5 mM KCl). The excitation and emission spectra were recorded at 390 nm, and 495 nm, respectively; using BioTek Synery Mx Monochromator based multi-mode plate reader. The increased acrylodan-Aha1 signals at 495 nm upon binding to Hsp90β was used to determine the dissociation constant (Kd) of the Aha1 and Hsp90β interaction using the equation:

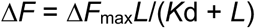

Where ΔF is change in the fluorescence intensity of the acrylodan-Aha1 upon binding to Hsp90β. ΔF_max_ is the maximum change in the fluorescence intensity of acrylodan-Aha1 when it is fully liganded with Hsp90β, and L is the concentration of Hsp90β. The ΔF_max_ value (50091 ± 4201) was calculated. ΔF was calculated by subtracting the fluorescence intensity of acrylodan-Aha1 in the absence of Hsp90β from the fluorescence intensity of acrylodan-Aha1 in the presence of Hsp90β ^123^. All the data were statistically analyzed by One-Way ANOVA and curve fitted using GraphPad software.

### NMR Spectroscopy of Aha1

Isotopically-labeled samples of Aha1, Aha1(1-163) and Aha1(164-338) were prepared by expression in *E. coli* BL21 DE3 [DNAY] in M9 minimal medium containing (^15^NH_4_)_2_SO_4_ and purified according to the protocol described above. Samples were dialyzed into NMR buffer (50 mM sodium phosphate buffer pH 7.00, 150 mM NaCl, 5 mM MgCl_2_, 2% glycerol, 2 mM DTT, 0.05% sodium azide). ^1^H-^15^N HSQC-TROSY spectra were acquired at 900 MHz on a Bruker Avance 900 spectrometer at 20ºC.

### Measurement of the effect SEW04874 on the Hsp90-Aha1 dependent refolding of luciferase

. Thermal inactivation ^124^ was used to study the refolding of luciferase with modifications. Firefly luciferase (0.1 µM; Promega) was thermally inactivated at 45°C for 8 min in 20 mM Hepes, pH 7.6, 85 mM potassium acetate, 1 mM magnesium acetate, 1.5 mM DDT and 1 mg/ml BSA containing 10 µM human Hsp90β. The thermally inactivated luciferase was diluted 10-fold in tubes containing the indicated molecular chaperones and co-chaperones, in the presence or absence of SEW04784 or geldanamycin (GA). Reaction mixtures were incubated at 24°C for 8 h to allow for luciferase refolding. Luciferase activity was measured using Promega Steady-Glo^®^ luciferase assay system and BioTek Synergy Mx Monochromator based multi-mode plate reader. The control was taken as 100 percent native firefly luciferase.

### *In vivo* GR and AR-mediated transcription assays

MDA-kb2 cells, which stably express an androgen- and glucocorticoid-responsive firefly luciferase reporter construct, were seeded at 20,000 cells/ well in 100 µl Leibovitz’s L-15 medium supplemented with 10% charcoal-stripped FBS at 0% CO_2,_ in 96 well clear bottom plates and grown to 80% confluence. Cells were treated with 0.2 nM DHT (EC_50_ for DHT in this cell line) or 10 nM dexamethasone (DEX; EC_50_ for DEX in this cell line) and the indicated concentrations of inhibitors for 20 h. Following treatment, the cells were lysed in 100 µl of Bright-Glo Luciferase Assay System (Promega Corp., Madison, WI) and incubated at room temperature for 4 min. For luciferase activity, the light emission was measure in a luminescence plate reader (Luminoskan Ascent, Thermo Labsystems) and luminescence measurement using Bright-Glo (Promega) luciferase assay reagent according to the manufacturer’s instructions.

### ELISA assays for prostate specific antigen secretion

ELISA assays to assess drug effects on PSA expression and secretion were performed in LNCaP and 22Rv1 prostate cancer cells. LNCaP cells endogenously express the AR T877A mutant (homozygous) and are sensitive to androgens. 22Rv1 cells express both full-length AR with an in-frame tandem duplication of exon 3 that encodes the second zinc finger of the AR DNA-binding domain, and a truncated AR lacking the C-terminal hormone binding domain. Thus, 22Rv1 cells are both androgen sensitive and can grow in the absence of androgens. Both LNCaP and 22Rv1 cells were maintained in MEM-EBSS (HyClone) supplemented with 10% FBS. The Human PSA ELISA Kit (Alpha Diagnostic International, ADI) to quantify secreted PSA in the media was used. The cells were plated at a density of 2 × 10^5^ cells per well in 12-well plates in 1 mL of respective standard growth media and incubated at 37° C with supplemental 5% CO_2_. When cells were attached, they were washed three times and replaced with 1 mL of respective growth media modified by replacement of 10% FBS with 10% CS-FBS at 37°C with supplemental 5% CO_2_. After 48 h, the cells were treated with DMSO + EtOH (negative control), DMSO + agonist (DHT, 50 nM for LNCaP and 1 nM for 22Rv1, positive control), or the agonist with a range of SEW concentrations for 24 h. For evaluating basal and hormone-induced PSA secretion in 22Rv1 cells, the aforementioned experimental conditions were used with the exception that SEW was tested at a single high dose (100 µM). Supernatants were collected, and the assays were performed according to manufacturer’s instructions. 25 µL of standards, controls, and supernatant samples were added into appropriate wells containing 100 µL of assay buffer in duplicate, covered, and incubated on a plate shaker (approximately at 200 rpm) for 60 min at room temperature. The wells were washed 3 times with 300 µL of 1X wash buffer followed by the addition of 100 µL of Ab-enzyme conjugate into each well, mixed gently for 10 secs, covered, and incubated the plate on a plate shaker (approximately at 200 rpm) for 30 min at room temperature. The wells were then washed 3 times with 300 µL of 1X wash buffer followed by the addition of 100 µL of TMB substrate per well, mixed gently for 10 secs, cover and incubated the plate on a plate shaker (approximately at 200 rpm) at room temperature until the samples turned dark blue. The reaction was stopped by adding 50 µL of stopping solution to all wells and mixed gently. The absorbance was measured at 450 nm with SpectraMAX 190 (Molecular Devices).

### GFP-tau assay

HEK293 cells stably carrying Dox-inducible GFP-tau (WT or P301L, 0N4R) were cultured in DMEM supplemented with 10% FBS and 1% P/S (Invitrogen). Cells were plated in 50 µL DMEM in 384-well plates (Greiner) coated with poly-D-lysine (Sigma). The next day, GFP-tau expression was induced by the addition of Dox (10 ng/mL). Compounds were then treated in 8-point dose starting at 20 µM in duplicate and incubated at 37 °C for 24 h. 30 minutes before the end of the 24-h incubation, cells were treated with Hoechst 33342 at the final concentration of 0.5 µg/ml. High-throughput imaging at 20X was performed on an InCell6000 automated microscope (GE) and GFP intensity of each cell was quantify using the InCell Analyzer Workstation (GE). To quantify effects of compounds tested, the median value of GFP intensity per cell of each well was computed, and normalized to the average median GFP intensity of all DMSO treated wells (32 wells in each plate). Cell count of each well was measured and normalized to the average cell count of all DMSO treated wells. Wells with a normalized cell count lower than 0.65 were disregarded. Curve fitting of the dose curve was conducted by Prism 7 (GraphPad).

### Brain slice quantitation of pS396/S404 tau

Quantitation performed as described^125^ with minor modifications.

## Acknowledgements

Support was provided by the National Institutes of Health (NIH) Grants HL095524, DK051870 and AG049665 to WEB. M.B.C., J.C.S and N.C.G. were partially supported by NIH grant 2G12MD007592 to the Border Biomedical Research Center (BBRC), from the National Institutes on Minority Health and Health Disparities (NIMHD). M.B.C is also supported by the Department of Defense (DOD) Prostate Cancer Research Program (PCRP) through grant number W81XWH-17-1-0435. Support was provided by the NIH Grant R01NS059690 to J.E.G. Support was provided by the NIH Grants GM057374 and GM113251 to J.H.D.

## References Cited

1 Karras, G. I. et al. HSP90 Shapes the Consequences of Human Genetic Variation. Cell 168, 856–866 e812, doi:10.1016/j.cell.2017.01.023 (2017).

2 Taipale, M. et al. A quantitative chaperone interaction network reveals the architecture of cellular protein homeostasis pathways. Cell 158, 434–448, doi:10.1016/j.cell.2014.05.039 (2014).

3 Rutz, D. A. et al. A switch point in the molecular chaperone Hsp90 responding to client interaction. Nat Commun 9, 1472, doi:10.1038/s41467-018-03946-x (2018).

4 Sahasrabudhe, P., Rohrberg, J., Biebl, M. M., Rutz, D. A. & Buchner, J. The Plasticity of the Hsp90 Co-chaperone System. Mol Cell 67, 947–961 e945, doi:10.1016/j.molcel.2017.08.004 (2017).

5 Schopf, F. H., Biebl, M. M. & Buchner, J. The HSP90 chaperone machinery. Nat Rev Mol Cell Biol 18, 345–360, doi:10.1038/nrm.2017.20 (2017).

6 Geller, R., Pechmann, S., Acevedo, A., Andino, R. & Frydman, J. Hsp90 shapes protein and RNA evolution to balance trade-offs between protein stability and aggregation. Nat Commun 9, 1781, doi:10.1038/s41467-018-04203-x (2018).

7 Powers, E. T. & Balch, W. E. Diversity in the origins of proteostasis networks--a driver for protein function in evolution. Nat Rev Mol Cell Biol 14, 237–248, doi:10.1038/nrm3542 (2013).

8 Balch, W. E., Morimoto, R. I., Dillin, A. & Kelly, J. W. Adapting proteostasis for disease intervention. Science 319, 916–919, doi:10.1126/science.1141448 (2008).

9 Labbadia, J. & Morimoto, R. I. The biology of proteostasis in aging and disease. Annu Rev Biochem 84, 435–464, doi:10.1146/annurev-biochem-060614-033955 (2015).

10 Li, J., Labbadia, J. & Morimoto, R. I. Rethinking HSF1 in Stress, Development, and Organismal Health. Trends Cell Biol 27, 895–905, doi:10.1016/j.tcb.2017.08.002 (2017).

11 Queitsch, C., Sangster, T. A. & Lindquist, S. Hsp90 as a capacitor of phenotypic variation. Nature 417, 618–624, doi:10.1038/nature749 (2002).

12 Balchin, D., Hayer-Hartl, M. & Hartl, F. U. In vivo aspects of protein folding and quality control. Science 353, aac4354, doi:10.1126/science.aac4354 (2016).

13 Sala, A. J., Bott, L. C. & Morimoto, R. I. Shaping proteostasis at the cellular, tissue, and organismal level. J Cell Biol 216, 1231–1241, doi:10.1083/jcb.201612111 (2017).

14 Verba, K. A. & Agard, D. A. How Hsp90 and Cdc37 Lubricate Kinase Molecular Switches. Trends Biochem Sci 42, 799–811, doi:10.1016/j.tibs.2017.07.002 (2017).

15 Frakes, A. E. & Dillin, A. The UPR(ER): Sensor and Coordinator of Organismal Homeostasis. Mol Cell 66, 761–771, doi:10.1016/j.molcel.2017.05.031 (2017).

16 Wang, C. & Balch, W. E. Bridging Genomics to Phenomics at Atomic Resolution through Variation Spatial Profiling. Cell Rep 24, 2013–2028 e2016, doi:10.1016/j.celrep.2018.07.059 (2018).

17 Stiegler, S. C. et al. A chemical compound inhibiting the Aha1-Hsp90 chaperone complex. J Biol Chem 292, 17073–17083, doi:10.1074/jbc.M117.797829 (2017).

18 Shelton, L. B. et al. Hsp90 activator Aha1 drives production of pathological tau aggregates. Proc Natl Acad Sci U S A 114, 9707–9712, doi:10.1073/pnas.1707039114 (2017).

19 Borges, J. C., Seraphim, T. V., Dores-Silva, P. R. & Barbosa, L. R. S. A review of multi-domain and flexible molecular chaperones studies by small-angle X-ray scattering. Biophys Rev 8, 107–120, doi:10.1007/s12551-016-0194-x (2016).

20 Ihrig, V. & Obermann, W. M. J. Identifying Inhibitors of the Hsp90-Aha1 Protein Complex, a Potential Target to Drug Cystic Fibrosis, by Alpha Technology. SLAS Discov 22, 923–928, doi:10.1177/2472555216688312 (2017).

21 Wolmarans, A., Lee, B., Spyracopoulos, L. & LaPointe, P. The Mechanism of Hsp90 ATPase Stimulation by Aha1. Sci Rep 6, 33179, doi:10.1038/srep33179 (2016).

22 Lepvrier, E. et al. Hsp90 Oligomers Interacting with the Aha1 Cochaperone: An Outlook for the Hsp90 Chaperone Machineries. Anal Chem 87, 7043–7051, doi:10.1021/acs.analchem.5b00051 (2015).

23 Rehn, A. B. & Buchner, J. p23 and Aha1. Subcell Biochem 78, 113–131, doi:10.1007/978-3-319-11731-7_6 (2015).

24 Synoradzki, K. & Bieganowski, P. Middle domain of human Hsp90 isoforms differentially binds Aha1 in human cells and alters Hsp90 activity in yeast. Biochim Biophys Acta 1853, 445–452, doi:10.1016/j.bbamcr.2014.11.026 (2015).

25 Tripathi, V., Darnauer, S., Hartwig, N. R. & Obermann, W. M. Aha1 can act as an autonomous chaperone to prevent aggregation of stressed proteins. J Biol Chem 289, 36220–36228, doi:10.1074/jbc.M114.590141 (2014).

26 Horvat, N. K. et al. A mutation in the catalytic loop of Hsp90 specifically impairs ATPase stimulation by Aha1p, but not Hch1p. J Mol Biol 426, 2379–2392, doi:10.1016/j.jmb.2014.04.002 (2014).

27 Calderwood, S. K. Molecular cochaperones: tumor growth and cancer treatment. Scientifica (Cairo) 2013, 217513, doi:10.1155/2013/217513 (2013).

28 Li, J. & Buchner, J. Structure, function and regulation of the hsp90 machinery. Biomed J 36, 106–117, doi:10.4103/2319-4170.113230 (2013).

29 Desjardins, F., Delisle, C. & Gratton, J. P. Modulation of the cochaperone AHA1 regulates heat-shock protein 90 and endothelial NO synthase activation by vascular endothelial growth factor. Arterioscler Thromb Vasc Biol 32, 2484–2492, doi:10.1161/ATVBAHA.112.256008 (2012).

30 Sun, L., Prince, T., Manjarrez, J. R., Scroggins, B. T. & Matts, R. L. Characterization of the interaction of Aha1 with components of the Hsp90 chaperone machine and client proteins. Biochim Biophys Acta 1823, 1092–1101, doi:10.1016/j.bbamcr.2012.03.014 (2012).

31 Holmes, J. L., Sharp, S. Y., Hobbs, S. & Workman, P. Silencing of HSP90 cochaperone AHA1 expression decreases client protein activation and increases cellular sensitivity to the HSP90 inhibitor 17-allylamino-17-demethoxygeldanamycin. Cancer Res 68, 1188–1197, doi:10.1158/0008-5472.CAN-07-3268 (2008).

32 Wang, X. et al. Hsp90 cochaperone Aha1 downregulation rescues misfolding of CFTR in cystic fibrosis. Cell 127, 803–815, doi:10.1016/j.cell.2006.09.043 (2006).

33 Li, J., Soroka, J. & Buchner, J. The Hsp90 chaperone machinery: conformational dynamics and regulation by co-chaperones. Biochim Biophys Acta 1823, 624–635, doi:10.1016/j.bbamcr.2011.09.003 (2012).

34 Meyer, P. et al. Structural basis for recruitment of the ATPase activator Aha1 to the Hsp90 chaperone machinery. EMBO J 23, 1402–1410, doi:10.1038/sj.emboj.7600141 (2004).

35 Wortmann, P., Gotz, M. & Hugel, T. Cooperative Nucleotide Binding in Hsp90 and Its Regulation by Aha1. Biophys J 113, 1711–1718, doi:10.1016/j.bpj.2017.08.032 (2017).

36 Taipale, M., Jarosz, D. F. & Lindquist, S. HSP90 at the hub of protein homeostasis: emerging mechanistic insights. Nat Rev Mol Cell Biol 11, 515–528, doi:10.1038/nrm2918 (2010).

37 Luo, Q., Boczek, E. E., Wang, Q., Buchner, J. & Kaila, V. R. Hsp90 dependence of a kinase is determined by its conformational landscape. Sci Rep 7, 43996, doi:10.1038/srep43996 (2017).

38 Koulov, A. V. et al. Biological and structural basis for Aha1 regulation of Hsp90 ATPase activity in maintaining proteostasis in the human disease cystic fibrosis. Mol Biol Cell 21, 871–884, doi:10.1091/mbc.E09-12-1017 (2010).

39 Li, J., Richter, K., Reinstein, J. & Buchner, J. Integration of the accelerator Aha1 in the Hsp90 co-chaperone cycle. Nat Struct Mol Biol 20, 326–331, doi:10.1038/nsmb.2502 (2013).

40 Retzlaff, M. et al. Asymmetric activation of the hsp90 dimer by its cochaperone aha1. Mol Cell 37, 344–354, doi:10.1016/j.molcel.2010.01.006 (2010).

41 Calamini, B. et al. Small-molecule proteostasis regulators for protein conformational diseases. Nat Chem Biol 8, 185–196, doi:10.1038/nchembio.763 (2011).

42 Coppinger, J. A. et al. A chaperone trap contributes to the onset of cystic fibrosis. PLoS One 7, e37682, doi:10.1371/journal.pone.0037682 (2012).

43 Khandelwal, A., Crowley, V. M. & Blagg, B. S. J. Natural Product Inspired N-Terminal Hsp90 Inhibitors: From Bench to Bedside? Med Res Rev 36, 92–118, doi:10.1002/med.21351 (2016).

44 Khandelwal, A. et al. Structure-guided design of an Hsp90beta N-terminal isoform-selective inhibitor. Nat Commun 9, 425, doi:10.1038/s41467-017-02013-1 (2018).

45 Roe, S. M. et al. Structural basis for inhibition of the Hsp90 molecular chaperone by the antitumor antibiotics radicicol and geldanamycin. J Med Chem 42, 260–266, doi:10.1021/jm980403y (1999).

46 Schulte, T. W. et al. Interaction of radicicol with members of the heat shock protein 90 family of molecular chaperones. Mol Endocrinol 13, 1435–1448, doi:10.1210/mend.13.9.0339 (1999).

47 Schulte, T. W. et al. Antibiotic radicicol binds to the N-terminal domain of Hsp90 and shares important biologic activities with geldanamycin. Cell Stress Chaperones 3, 100– 108 (1998).

48 Sharma, S. V., Agatsuma, T. & Nakano, H. Targeting of the protein chaperone, HSP90, by the transformation suppressing agent, radicicol. Oncogene 16, 2639–2645, doi:10.1038/sj.onc.1201790 (1998).

49 Taldone, T., Ochiana, S. O., Patel, P. D. & Chiosis, G. Selective targeting of the stress chaperome as a therapeutic strategy. Trends Pharmacol Sci 35, 592–603, doi:10.1016/j.tips.2014.09.001 (2014).

50 Whitesell, L., Mimnaugh, E. G., De Costa, B., Myers, C. E. & Neckers, L. M. Inhibition of heat shock protein HSP90-pp60v-src heteroprotein complex formation by benzoquinone ansamycins: essential role for stress proteins in oncogenic transformation. Proc Natl Acad Sci U S A 91, 8324–8328 (1994).

51 Zhang, Z., You, Z., Dobrowsky, R. T. & Blagg, B. S. J. Synthesis and evaluation of a ring-constrained Hsp90 C-terminal inhibitor that exhibits neuroprotective activity. Bioorg Med Chem Lett, doi:10.1016/j.bmcl.2018.03.071 (2018).

52 Taldone, T., Patel, H. J., Bolaender, A., Patel, M. R. & Chiosis, G. Protein chaperones: a composition of matter review (2008 - 2013). Expert Opin Ther Pat 24, 501–518, doi:10.1517/13543776.2014.887681 (2014).

53 Shrestha, L., Patel, H. J. & Chiosis, G. Chemical Tools to Investigate Mechanisms Associated with HSP90 and HSP70 in Disease. Cell Chem Biol 23, 158–172, doi:10.1016/j.chembiol.2015.12.006 (2016).

54 Burlison, J. A. & Blagg, B. S. Synthesis and evaluation of coumermycin A1 analogues that inhibit the Hsp90 protein folding machinery. Org Lett 8, 4855–4858, doi:10.1021/ol061918j (2006).

55 Eccles, S. A. et al. NVP-AUY922: a novel heat shock protein 90 inhibitor active against xenograft tumor growth, angiogenesis, and metastasis. Cancer Res 68, 2850–2860, doi:10.1158/0008-5472.CAN-07-5256 (2008).

56 Lin, T. Y. et al. The novel HSP90 inhibitor STA-9090 exhibits activity against Kit-dependent and-independent malignant mast cell tumors. Exp Hematol 36, 1266–1277, doi:10.1016/j.exphem.2008.05.001 (2008).

57 Neckers, L. & Workman, P. Hsp90 molecular chaperone inhibitors: are we there yet? Clin Cancer Res 18, 64–76, doi:10.1158/1078-0432.CCR-11-1000 (2012).

58 Glaze, E. R. et al. Preclinical toxicity of a geldanamycin analog, 17-(dimethylaminoethylamino)-17-demethoxygeldanamycin (17-DMAG), in rats and dogs: potential clinical relevance. Cancer Chemother Pharmacol 56, 637–647, doi:10.1007/s00280-005-1000-9 (2005).

59 Iyer, G. et al. A phase I trial of docetaxel and pulse-dose 17-allylamino-17-demethoxygeldanamycin in adult patients with solid tumors. Cancer Chemother Pharmacol 69, 1089–1097, doi:10.1007/s00280-011-1789-3 (2012).

60 Shrestha, L., Bolaender, A., Patel, H. J. & Taldone, T. Heat Shock Protein (HSP) Drug Discovery and Development: Targeting Heat Shock Proteins in Disease. Curr Top Med Chem 16, 2753–2764 (2016).

61 Butler, L. M., Ferraldeschi, R., Armstrong, H. K., Centenera, M. M. & Workman, P. Maximizing the Therapeutic Potential of HSP90 Inhibitors. Mol Cancer Res 13, 1445–1451, doi:10.1158/1541-7786.MCR-15-0234 (2015).

62 Wang, M. et al. Development of Heat Shock Protein (Hsp90) Inhibitors To Combat Resistance to Tyrosine Kinase Inhibitors through Hsp90-Kinase Interactions. J Med Chem 59, 5563–5586, doi:10.1021/acs.jmedchem.5b01106 (2016).

63 Cohen, S. M. et al. Novel C-terminal Hsp90 inhibitor for head and neck squamous cell cancer (HNSCC) with in vivo efficacy and improved toxicity profiles compared with standard agents. Ann Surg Oncol 19 Suppl 3, S483–490, doi:10.1245/s10434-011-1971-1 (2012).

64 Hall, J. A. et al. Novobiocin Analogues That Inhibit the MAPK Pathway. J Med Chem 59, 925–933, doi:10.1021/acs.jmedchem.5b01354 (2016).

65 Kusuma, B. R. et al. Synthesis and evaluation of novologues as C-terminal Hsp90 inhibitors with cytoprotective activity against sensory neuron glucotoxicity. J Med Chem 55, 5797–5812, doi:10.1021/jm300544c (2012).

66 Liu, W. et al. KU675, a Concomitant Heat-Shock Protein Inhibitor of Hsp90 and Hsc70 that Manifests Isoform Selectivity for Hsp90alpha in Prostate Cancer Cells. Mol Pharmacol 88, 121–130, doi:10.1124/mol.114.097303 (2015).

67 Samadi, A. K. et al. A novel C-terminal HSP90 inhibitor KU135 induces apoptosis and cell cycle arrest in melanoma cells. Cancer Lett 312, 158–167, doi:10.1016/j.canlet.2011.07.031 (2011).

68 Zhang, L., Zhao, H., Blagg, B. S. & Dobrowsky, R. T. C-terminal heat shock protein 90 inhibitor decreases hyperglycemia-induced oxidative stress and improves mitochondrial bioenergetics in sensory neurons. J Proteome Res 11, 2581–2593, doi:10.1021/pr300056m (2012).

69 Zhao, H. & Blagg, B. S. Novobiocin analogues with second-generation noviose surrogates. Bioorg Med Chem Lett 23, 552–557, doi:10.1016/j.bmcl.2012.11.022 (2013).

70 Zuck, P. et al. Miniaturization of absorbance assays using the fluorescent properties of white microplates. Anal Biochem 342, 254–259, doi:10.1016/j.ab.2005.04.029 (2005).

71 Moses, M. A. et al. Targeting the Hsp40/Hsp70 Chaperone Axis as a Novel Strategy to Treat Castration-Resistant Prostate Cancer. Cancer Res 78, 4022–4035, doi:10.1158/0008-5472.CAN-17-3728 (2018).

72 Wegele, H., Wandinger, S. K., Schmid, A. B., Reinstein, J. & Buchner, J. Substrate transfer from the chaperone Hsp70 to Hsp90. J Mol Biol 356, 802–811, doi:10.1016/j.jmb.2005.12.008 (2006).

73 Li, X. et al. Analogs of the Allosteric Heat Shock Protein 70 (Hsp70) Inhibitor, MKT-077, as Anti-Cancer Agents. ACS Med Chem Lett 4, doi:10.1021/ml400204n (2013).

74 Gupta, R. et al. Firefly luciferase mutants as sensors of proteome stress. Nat Methods 8, 879–884, doi:10.1038/nmeth.1697 (2011).

75 De Leon, J. T. et al. Targeting the regulation of androgen receptor signaling by the heat shock protein 90 cochaperone FKBP52 in prostate cancer cells. Proc Natl Acad Sci U S A 108, 11878–11883, doi:10.1073/pnas.1105160108 (2011).

76 Elo, J. P. et al. Mutated human androgen receptor gene detected in a prostatic cancer patient is also activated by estradiol. J Clin Endocrinol Metab 80, 3494–3500, doi:10.1210/jcem.80.12.8530589 (1995).

77 Grigoryev, D. N., Long, B. J., Njar, V. C. & Brodie, A. H. Pregnenolone stimulates LNCaP prostate cancer cell growth via the mutated androgen receptor. J Steroid Biochem Mol Biol 75, 1–10 (2000).

78 Long, B. J. et al. Antiandrogenic effects of novel androgen synthesis inhibitors on hormone-dependent prostate cancer. Cancer Res 60, 6630–6640 (2000).

79 Miyamoto, H., Yeh, S., Wilding, G. & Chang, C. Promotion of agonist activity of antiandrogens by the androgen receptor coactivator, ARA70, in human prostate cancer DU145 cells. Proc Natl Acad Sci U S A 95, 7379–7384 (1998).

80 Tan, J. et al. Dehydroepiandrosterone activates mutant androgen receptors expressed in the androgen-dependent human prostate cancer xenograft CWR22 and LNCaP cells. Mol Endocrinol 11, 450–459, doi:10.1210/mend.11.4.9906 (1997).

81 Yeh, S., Miyamoto, H., Shima, H. & Chang, C. From estrogen to androgen receptor: a new pathway for sex hormones in prostate. Proc Natl Acad Sci U S A 95, 5527–5532 (1998).

82 Drubin, D. G. & Kirschner, M. W. Tau protein function in living cells. J Cell Biol 103, 2739–2746 (1986).

83 Weingarten, M. D., Lockwood, A. H., Hwo, S. Y. & Kirschner, M. W. A protein factor essential for microtubule assembly. Proc Natl Acad Sci U S A 72, 1858–1862 (1975).

84 Hong, M. et al. Mutation-specific functional impairments in distinct tau isoforms of hereditary FTDP-17. Science 282, 1914–1917 (1998).

85 Hutton, M. et al. Association of missense and 5’-splice-site mutations in tau with the inherited dementia FTDP-17. Nature 393, 702–705, doi:10.1038/31508 (1998).

86 von Bergen, M. et al. Assembly of tau protein into Alzheimer paired helical filaments depends on a local sequence motif ((306)VQIVYK(311)) forming beta structure. Proc Natl Acad Sci U S A 97, 5129–5134 (2000).

87 Young, Z. T., Mok, S. A. & Gestwicki, J. E. Therapeutic Strategies for Restoring Tau Homeostasis. Cold Spring Harb Perspect Med 8, doi:10.1101/cshperspect.a024612 (2018).

88 Shelton, L. B., Koren, J., 3rd & Blair, L. J. Imbalances in the Hsp90 Chaperone Machinery: Implications for Tauopathies. Front Neurosci 11, 724, doi:10.3389/fnins.2017.00724 (2017).

89 Rauch, J. N., Olson, S. H. & Gestwicki, J. E. Interactions between Microtubule-Associated Protein Tau (MAPT) and Small Molecules. Cold Spring Harb Perspect Med 7, doi:10.1101/cshperspect.a024034 (2017).

90 Pratt, W. B., Gestwicki, J. E., Osawa, Y. & Lieberman, A. P. Targeting Hsp90/Hsp70-based protein quality control for treatment of adult onset neurodegenerative diseases. Annu Rev Pharmacol Toxicol 55, 353–371, doi:10.1146/annurev-pharmtox-010814-124332 (2015).

91 Spillantini, M. G. & Goedert, M. Tau pathology and neurodegeneration. Lancet Neurol 12, 609–622, doi:10.1016/S1474-4422(13)70090-5 (2013).

92 Dickey, C. A. et al. HSP induction mediates selective clearance of tau phosphorylated at proline-directed Ser/Thr sites but not KXGS (MARK) sites. FASEB J 20, 753–755, doi:10.1096/fj.05-5343fje (2006).

93 Dickey, C. A. et al. The high-affinity HSP90-CHIP complex recognizes and selectively degrades phosphorylated tau client proteins. J Clin Invest 117, 648–658, doi:10.1172/JCI29715 (2007).

94 Dickey, C. A. et al. Akt and CHIP coregulate tau degradation through coordinated interactions. Proc Natl Acad Sci U S A 105, 3622–3627, doi:10.1073/pnas.0709180105 (2008).

95 Jinwal, U. K. et al. Chemical manipulation of hsp70 ATPase activity regulates tau stability. J Neurosci 29, 12079–12088, doi:10.1523/JNEUROSCI.3345-09.2009 (2009).

96 Luo, W. et al. Roles of heat-shock protein 90 in maintaining and facilitating the neurodegenerative phenotype in tauopathies. Proc Natl Acad Sci U S A 104, 9511–9516, doi:10.1073/pnas.0701055104 (2007).

97 O’Leary, J. C., 3rd et al. Phenothiazine-mediated rescue of cognition in tau transgenic mice requires neuroprotection and reduced soluble tau burden. Mol Neurodegener 5, 45, doi:10.1186/1750-1326-5-45 (2010).

98 Jinwal, U. K. et al. Cdc37/Hsp90 protein complex disruption triggers an autophagic clearance cascade for TDP-43 protein. J Biol Chem 287, 24814–24820, doi:10.1074/jbc.M112.367268 (2012).

99 Kinoshita, E. et al. Profiling of protein thiophosphorylation by Phos-tag affinity electrophoresis: evaluation of adenosine 5’-O-(3-thiotriphosphate) as a phosphoryl donor in protein kinase reactions. Proteomics 14, 668–679, doi:10.1002/pmic.201300533 (2014).

100 Guthrie, C. R. & Kraemer, B. C. Proteasome inhibition drives HDAC6-dependent recruitment of tau to aggresomes. J Mol Neurosci 45, 32–41, doi:10.1007/s12031-011-9502-x (2011).

101 Hasegawa, M., Smith, M. J. & Goedert, M. Tau proteins with FTDP-17 mutations have a reduced ability to promote microtubule assembly. FEBS Lett 437, 207–210 (1998).

102 Pratt, W. B. & Dittmar, K. D. Studies with Purified Chaperones Advance the Understanding of the Mechanism of Glucocorticoid Receptor-hsp90 Heterocomplex Assembly. Trends Endocrinol Metab 9, 244–252 (1998).

103 Pratt, W. B., Morishima, Y., Gestwicki, J. E., Lieberman, A. P. & Osawa, Y. A model in which heat shock protein 90 targets protein-folding clefts: rationale for a new approach to neuroprotective treatment of protein folding diseases. Exp Biol Med (Maywood) 239, 1405–1413, doi:10.1177/1535370214539444 (2014).

104 Attard, G. et al. Phase I clinical trial of a selective inhibitor of CYP17, abiraterone acetate, confirms that castration-resistant prostate cancer commonly remains hormone driven. J Clin Oncol 26, 4563–4571, doi:10.1200/JCO.2007.15.9749 (2008).

105 Tran, C. et al. Development of a second-generation antiandrogen for treatment of advanced prostate cancer. Science 324, 787–790, doi:10.1126/science.1168175 (2009).

106 Antonarakis, E. S. et al. AR-V7 and resistance to enzalutamide and abiraterone in prostate cancer. N Engl J Med 371, 1028–1038, doi:10.1056/NEJMoa1315815 (2014).

107 Arora, V. K. et al. Glucocorticoid receptor confers resistance to antiandrogens by bypassing androgen receptor blockade. Cell 155, 1309–1322, doi:10.1016/j.cell.2013.11.012 (2013).

108 Sahu, B. et al. FoxA1 specifies unique androgen and glucocorticoid receptor binding events in prostate cancer cells. Cancer Res 73, 1570–1580, doi:10.1158/0008-5472.CAN-12-2350 (2013).

109 Szmulewitz, R. Z. et al. Serum/glucocorticoid-regulated kinase 1 expression in primary human prostate cancers. Prostate 72, 157–164, doi:10.1002/pros.21416 (2012).

110 Yemelyanov, A. et al. Differential targeting of androgen and glucocorticoid receptors induces ER stress and apoptosis in prostate cancer cells: a novel therapeutic modality. Cell Cycle 11, 395–406, doi:10.4161/cc.11.2.18945 (2012).

111 Gillis, J. L. et al. Constitutively-active androgen receptor variants function independently of the HSP90 chaperone but do not confer resistance to HSP90 inhibitors. Oncotarget 4, 691–704, doi:10.18632/oncotarget.975 (2013).

112 Shafi, A. A., Cox, M. B. & Weigel, N. L. Androgen receptor splice variants are resistant to inhibitors of Hsp90 and FKBP52, which alter androgen receptor activity and expression. Steroids 78, 548–554, doi:10.1016/j.steroids.2012.12.013 (2013).

113 Echeverria, P. C., Bernthaler, A., Dupuis, P., Mayer, B. & Picard, D. An interaction network predicted from public data as a discovery tool: application to the Hsp90 molecular chaperone machine. PLoS One 6, e26044, doi:10.1371/journal.pone.0026044 (2011).

114 Zierer, B. K. et al. Importance of cycle timing for the function of the molecular chaperone Hsp90. Nat Struct Mol Biol 23, 1020–1028, doi:10.1038/nsmb.3305 (2016).

115 Karagoz, G. E. et al. Hsp90-Tau complex reveals molecular basis for specificity in chaperone action. Cell 156, 963–974, doi:10.1016/j.cell.2014.01.037 (2014).

116 Mok, S. A. et al. Mapping interactions with the chaperone network reveals factors that protect against tau aggregation. Nat Struct Mol Biol 25, 384–393, doi:10.1038/s41594-018-0057-1 (2018).

117 Boselli, M. et al. An inhibitor of the proteasomal deubiquitinating enzyme USP14 induces tau elimination in cultured neurons. J Biol Chem 292, 19209–19225, doi:10.1074/jbc.M117.815126 (2017).

118 Chang, H. C. et al. Hsc70 is required for endocytosis and clathrin function in Drosophila. J Cell Biol 159, 477–487, doi:10.1083/jcb.200205086 (2002).

119 Rauch, J. N. & Gestwicki, J. E. Binding of human nucleotide exchange factors to heat shock protein 70 (Hsp70) generates functionally distinct complexes in vitro. J Biol Chem 289, 1402–1414, doi:10.1074/jbc.M113.521997 (2014).

120 Carrigan, P. E., Sikkink, L. A., Smith, D. F. & Ramirez-Alvarado, M. Domain:domain interactions within Hop, the Hsp70/Hsp90 organizing protein, are required for protein stability and structure. Protein Sci 15, 522–532, doi:10.1110/ps.051810106 (2006).

121 Weaver, A. J., Sullivan, W. P., Felts, S. J., Owen, B. A. & Toft, D. O. Crystal structure and activity of human p23, a heat shock protein 90 co-chaperone. J Biol Chem 275, 23045–23052, doi:10.1074/jbc.M003410200 (2000).

122 Chang, L. et al. High-throughput screen for small molecules that modulate the ATPase activity of the molecular chaperone DnaK. Anal Biochem 372, 167–176, doi:10.1016/j.ab.2007.08.020 (2008).

123 Singh, J. K., Makde, R. D., Kumar, V. & Panda, D. A membrane protein, EzrA, regulates assembly dynamics of FtsZ by interacting with the C-terminal tail of FtsZ. Biochemistry 46, 11013–11022, doi:10.1021/bi700710j (2007).

124 Minami, Y. & Minami, M. Hsc70/Hsp40 chaperone system mediates the Hsp90-dependent refolding of firefly luciferase. Genes Cells 4, 721–729 (1999).

125 Congdon, E. E. et al. Methylthioninium chloride (methylene blue) induces autophagy and attenuates tauopathy in vitro and in vivo. Autophagy 8, 609–622, doi:10.4161/auto.19048 (2012).

